# Fine-mapping *cis*-regulatory variants in diverse human populations

**DOI:** 10.1101/384396

**Authors:** Ashley K. Tehranchi, Brian Hie, Michael Dacre, Irene M. Kaplow, Kade P Pettie, Peter A. Combs, Hunter B. Fraser

## Abstract

Genome-wide association studies (GWAS) are a powerful approach for connecting genotype to phenotype. Most GWAS hits are located in cis-regulatory regions, but the underlying causal variants and their molecular mechanisms remain unknown. To better understand human *cis*-regulatory variation, we mapped quantitative trait loci for chromatin accessibility (caQTLs)—a key step in cis-regulation—in 1000 individuals from 10 diverse populations. Most caQTLs were shared across populations, allowing us to leverage the genetic diversity to fine-map candidate causal regulatory variants, several thousand of which have been previously implicated in GWAS. In addition, many caQTLs that affect the expression of distal genes also alter the landscape of long-range chromosomal interactions, suggesting a mechanism for long-range expression QTLs. In sum, our results show that molecular QTL mapping integrated across diverse populations provides a high-resolution view of how worldwide human genetic variation affects chromatin accessibility, gene expression, and phenotype.

## Introduction

GWAS have been instrumental in advancing our understanding of complex traits. Indeed, these studies have successfully mapped thousands of loci associated with hundreds of human diseases and other traits (Eicher et al. 2015). However, the vast majority of causal variants that drive the associations remain unknown, since GWAS resolution is limited by the correlations between nearby variants known as linkage disequilibrium (LD). Pinpointing causal variants is essential not only for accurately predicting disease risk, but also for understanding the molecular mechanisms that underlie complex trait variation. Increasing GWAS sample size or genotyping density can improve resolution, but even the largest studies have yielded only a few dozen causal variants (Farh et al. 2015, Huang et al. 2017).

An alternative to increasing sample size within a single population is trans-ethnic fine-mapping, in which a GWAS is performed across multiple populations (van de Bunt et al. 2015, Asimit et al. 2016). Shared causal variants should be consistently associated with a trait across populations, while tag SNPs—those associated only because of their LD with a causal variant—may only be associated in a subset of populations, due to differing LD structures. This approach is especially effective when combining populations with disparate LD patterns, such as Europeans and Africans (Asimit et al. 2016). However, most GWAS have been restricted to European cohorts, limiting not only their mapping resolution, but also their ability to predict disease risk in non-European individuals (Hindorff et al. 2018).

Another approach to understanding molecular mechanisms underlying complex traits has been to intersect GWAS hits with quantitative trait loci (QTLs) for molecular-level traits, which provide an effective “stepping stone” on the path from genotype to disease. QTLs involved in *cis*-regulation are especially informative, since the vast majority of GWAS hits are in regulatory regions (Hindorff et al., 2009). For example, QTLs for mRNA levels (eQTLs) are enriched for SNPs implicated by GWAS in a wide range of diseases, suggesting specific genes as likely mediators of the associations (Lappalainen et al. 2013, GTEx Consortium 2017). Similar enrichments have been reported for QTLs affecting mRNA splicing (Fraser and Xie 2009, Li et al 2016), transcription factor binding (Waszak et al. 2015, Tehranchi et al. 2016), and chromatin accessibility (Kumasaka et al. 2016, Gate et al. 2018). However, nearly all of these molecular-level QTL studies have been limited to a single population (most often either Europeans or Yorubans), thus limiting their utility for understanding worldwide human diversity—similar to th bias of GWAS for Europeans. Yet there is a great deal we could learn about the genetic basis of phenotypic diversity from performing trans-ethnic fine-mapping on molecular-level QTLs.

We recently developed an efficient approach for mapping molecular QTLs in which samples are pooled prior to sequencing. This pooling minimizes experimental variability between samples (both within and between experimental “batches”), reducing the cost and effort of QTL mapping by over 25-fold compared to standard unpooled approaches (Tehranchi et al. 2016). However, our initial study suffered from the same limitations as most other molecular QTL studies: it involved a small cohort (60 individuals) from a single population. We reasoned that the efficiency of pooling could enable us to map cis-regulatory QTLs with far greater numbers of individuals and populations than would otherwise be possible, allowing us to perform trans-ethnic fine-mapping to pinpoint thousands of likely causal variants affecting cis-regulation and, in many cases, complex disease risk.

## Results

### Pooled QTL mapping of variants affecting chromatin accessibility

We applied pooled QTL mapping (Kaplow et al. 2015, Tehranchi et al. 2016) to chromatin accessibility (CA), a reliable indicator of local cis-regulatory activity, as measured by the Assay for Transposase-Accessible Chromatin (ATAC-seq) (Buenrostro et al. 2015). In this approach, many samples are combined into a single pool, in which ATAC-seq (or another assay of interest) is performed only once. Genetic variants that affect CA in *cis* will cause the more accessible allele to increase in frequency after ATAC compared to before, whereas variants with no effect on accessibility will have no significant change in frequency (Figure 1A). For each SNP, we estimated the post-ATAC allele frequency from read counts of each allele present in the ATAC-seq reads and used a regression-based approach to estimate pre-ATAC allele frequency; the significance of the difference between these two frequencies is our caQTL p-value (Tehranchi et al. 2016).

**Figure 1.**
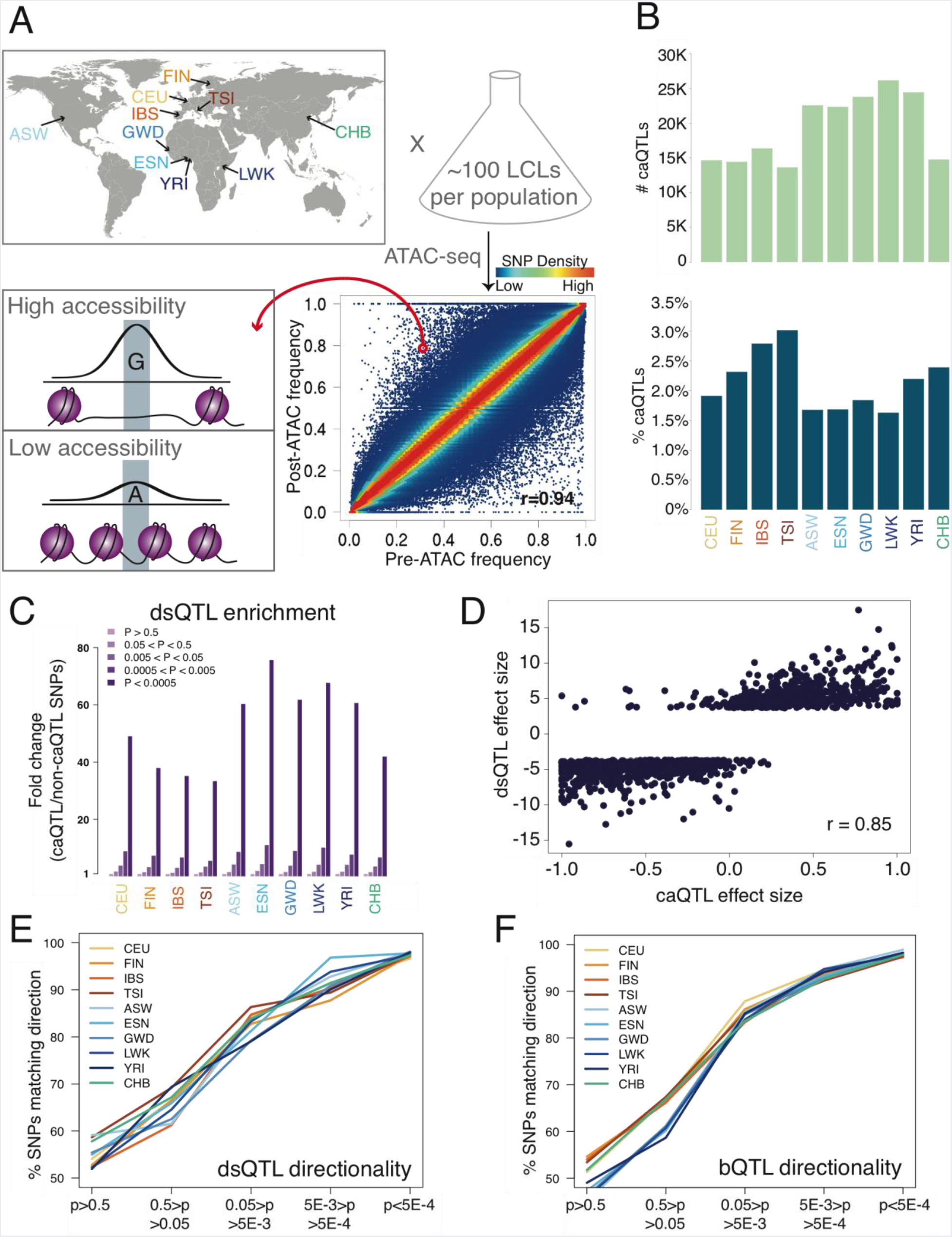
Outline and results of pooled ATAC-seq. A. Performing ATAC-seq in a pool of individuals selects DNA molecules with higher CA, thus enriching for more accessible alleles. In this example (ASW population), the G allele has a low pre-ATAC frequency but a high post-ATAC frequency, due to its increased CA. The ten population abbreviations refer to: CEU, Utah residents with North European ancestry; FIN, Finnish; TSI, Tuscan; IBS, Iberian; ASW, African-American from Southwest US; YRI, Yoruban; ESN, Esan; LWK, Luhya; GWD, Gambian; and CHB, Han Chinese. **B**. The number of caQTLs (top), and the percent of all tested SNPs called as caQTLs (bottom). **C**. Enrichment of caQTLs among dsQTLs (Degner et al. 2012), at a range of caQTL p-value cutoffs. **D**. Quantitative effect sizes of caQTLs and dsQTLs are highly correlated (scales of each axis are not comparable, and do not affect the correlation coefficient). **E-F**. The degree of allelic concordance between our caQTLs and: **E**. dsQTLs (Degner et al. 2012). **F**. bQTLs aggregated for five TFs (Tehranchi et al. 2016).

We selected a total of 1000 lymphoblastoid cell lines (LCLs) with full genome sequences (1000 Genomes Project Consortium 2015) from 10 diverse populations: four European, four African, one African-American, and one Han Chinese (Figure 1A, Supplementary file 1). We combined 66-112 unrelated individuals from each population into one pool per population, and performed ATAC-seq with two biological replicates per population (Figure 1--figure supplement 1), resulting in a total of ~808 million informative autosomal reads from 20 pools. All populations showed a high correlation between pre-ATAC and post-ATAC allele frequencies (average Pearson r = 0.92; Figure 1A, Figure 1--figure supplement 2), as expected if most SNPs do not affect CA and thus have similar pre- and post-ATAC frequencies (Tehranchi et al. 2016). At a nominal*p* < 5x10^-4^—corresponding to an irreproducible discovery rate (IDR, analogous to a false discovery rate; Figure 1--figure supplement 3) of ~1%—we mapped 13,657 to 26,182 independent caQTLs per population. This comprised 1.5%-3.0% of all testable SNPs, defined as those with >2% minor allele frequency (MAF) and covered by at least 20 reads ( Figure 1B, Figure 1--figure supplement 4). A total of 126,773 caQTLs were significant in at least one population (Supplementary file 2).

### Accuracy and efficiency of pooled QTL mapping

To gauge the accuracy of our caQTLs, we compared them with previously mapped QTLs. DNase-seq is another method used to assay chromatin accessibility, and SNPs affecting DNase read density—known as dsQTLs—have been mapped using 70 individual (non-pooled) Yoruban LCLs (Degner et al. 2012). We compared our caQTLs to dsQTLs in three ways. First, testing the overlap between these two sets, we found a 33- to 76-fold enrichment of dsQTLs among caQTLs, with highest enrichment in the five African/African-American populations ( Figure 1C; enrichments for other QTLs in Figure 1--figure supplement 5). Second, we compared the quantitative effect sizes of caQTLs and dsQTLs, and found excellent agreement (Pearson *r* = 0.85; Figure 1D). Third, we tested how the directionality agreement (whether caQTLs and dsQTLs call the same allele as more accessible) changes with the caQTL p-value, and found that agreement increased with more stringent cutoffs, reaching 96.7%-98.1% agreement at caQTL p < 5x10^-4^ ( Figure 1E). These three tests show that our caQTLs are consistent with dsQTLs mapped with unpooled samples.

Because accessible chromatin is more often bound by TFs, we also tested the directionality agreement between overlapping caQTLs and TF binding QTLs (bQTLs) aggregated for five TFs (Tehranchi et al. 2016), since we expect increased TF binding to be associated with increased CA. Once again, we found that the agreement increases with more stringent p-value cutoffs, reaching 97.3%-98.9% agreement at caQTL p < 5x10^-4^ ( Figure 1F). This high level of agreement suggests a low rate of false positive caQTLs, consistent with our estimated 1% IDR.

To assess the efficiency of our pooling approach, we compared our results with caQTLs mapped with RASQUAL—a computational QTL mapping approach that accounts for many possible confounding variables—in non-pooled European LCLs (Kumasaka et al. 2016). At a matched cutoff in our CEU population, we mapped 2.8-fold more caQTLs with 15-fold fewer reads and 12-fold fewer ATAC-seq libraries—a ~40-fold improvement in cost per caQTL (see Supplemental Note). Overall, our comparisons with previously mapped QTLs (Figure 1C-F, Figure 1--figure supplement 5) suggest that pooling is an efficient strategy that agrees well with unpooled QTL mapping.

### caQTLs shared across populations

We next assessed the extent of caQTL sharing across populations. A caQTL might not be shared due to biological causes, such as dependence of a variant’s effect on the genetic background (epistasis), or due to trivial causes such as not meeting our 20 read cutoff in one population. To exclude trivial cases we focused on 142,049 variants that were testable in all ten populations. Among these, we observed a clear trend for increased sharing within continents: the mean fraction of shared caQTLs was 59.9% within Africans and 59.8% within Europeans, compared to 48.4% between these two groups (Figure 2A). The Han Chinese (CHB) caQTLs were shared moderately with Europeans (52.4%) and less well with Africans (47.7%), reflecting their closer relatedness to Europeans. In addition, African-American (ASW) caQTLs showed greater sharing with all five European/Han populations than any of the four African populations did, consistent with their admixed ancestry. Restricting the analysis to caQTLs with similar allele frequencies across populations led to a similar pattern (Figure 2--figure supplement 1). The concordance between caQTL sharing and known phylogenetic relationships suggests that some caQTLs may have population-specific effects, as has also been observed for other types of molecular QTLs (Stranger et al. 2012, Fraser et al. 2012), though the biological mechanisms underlying this divergence will require further study.

**Figure 2.**
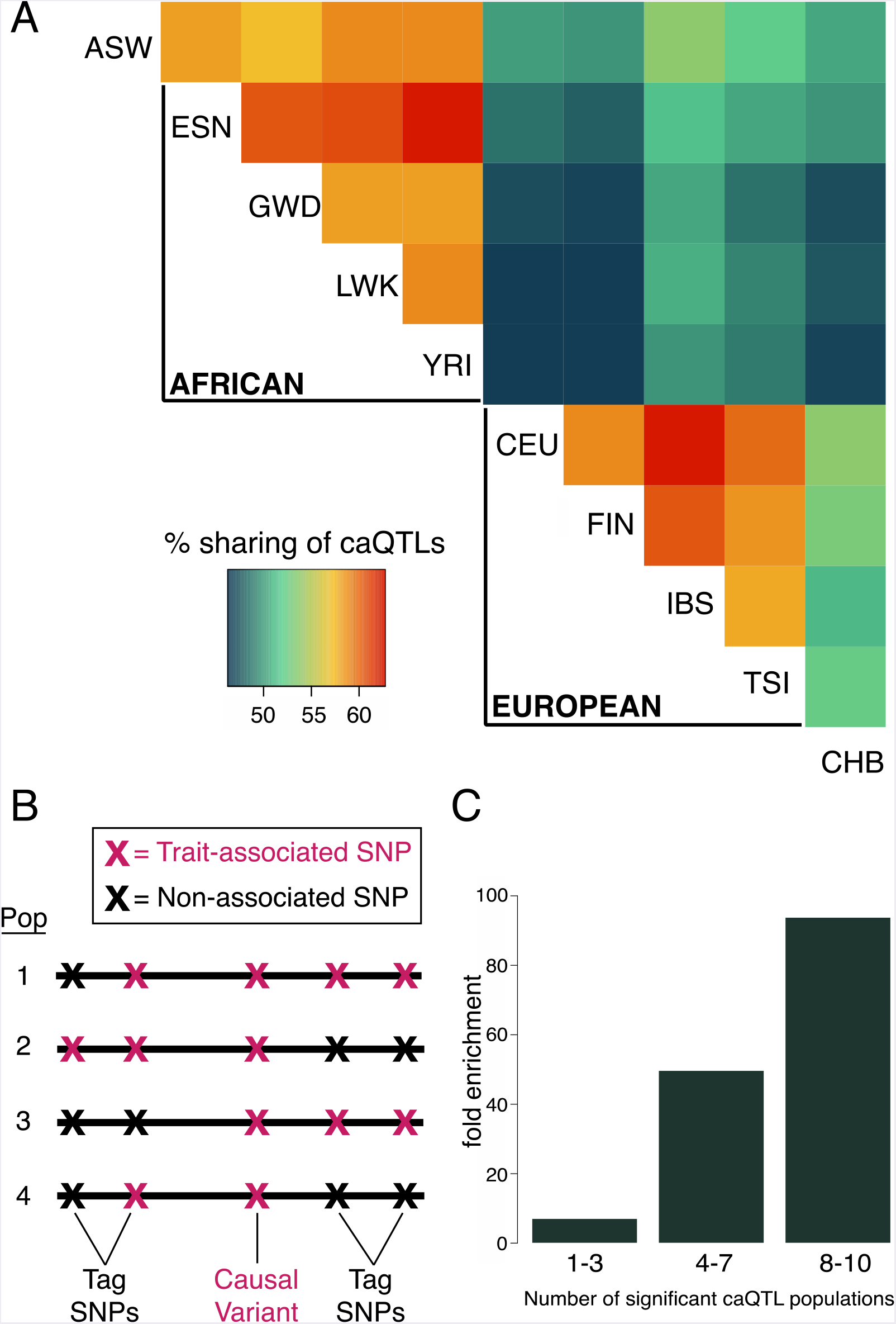
Fine-mapping shared caQTLs. A. Heatmap showing the overlap in caQTLs for every pair of populations (only for variants that were testable in all ten). To avoid issues related to arbitrary p-value cutoffs, we used the shift in p-value distribution, known as n_1_ (Storey et al. 2004), to assess overlap. **B**. Mapping a trait in multiple populations differing in LD structure allows fine-mapping of causal variants, which will show the most consistent associations. **C**. caQTLs shared across many populations (at p < 5x10^-4^) are more highly enriched for experimentally-determined causal eQTL variants (Tewhey et al. 2016).

A fundamental limitation of human GWAS is their inability to pinpoint most causal variants, due to linkage disequilibrium (LD) that often results in many variants with equally strong associations (van de Bunt et al. 2015). However, causal variants can be fine-mapped by combining association data across populations, especially with African populations due to their low LD (Asimit et al. 2016). Causal variants should be consistently associated with a trait across populations, while tag SNPs—those associated only because of their LD with a causal variant—may only be associated in a subset of populations, due to differing LD structures ( Figure 2B).

We performed fine-mapping by searching for caQTLs present in multiple populations. To test whether this was indeed enriching for causal variants, we intersected our caQTLs with a collection of experimentally verified causal eQTL variants (Tewhey et al. 2016). We found increasing enrichment for known causal variants with increasing number of significant caQTL populations ( Figure 2C), suggesting that combining the diverse LD patterns improved our mapping resolution substantially. In order to account for both the number of significant populations as well as the caQTL strength within each population (Figure 2--figure supplement 2), we calculated Fisher’s combined p-value for each caQTL across all ten populations; at a combined p < 5x10^-6^, we identified 45,243 SNPs, which we refer to as “shared caQTLs” (Supplementary file 3). Nearly all (99.8%) of these were significant across multiple populations, and 98.6% showed concordance of the more accessible allele across populations.

### Characterizing fine-mapped caQTLs

Leveraging the high resolution of our fine-mapped shared caQTLs, we first investigated their genomic locations. We found that they were most highly enriched near active enhancers and transcription start sites (TSSs), accounting for 54% of shared caQTLs in just 3.1% of the genome ( Figure 3A, purple and dark green). However, these enrichments were primarily driven by ATAC-seq read density, reflecting greater chromatin accessibility in these regions; after controlling for read density, we found the strongest enrichments in weak enhancers (2.3-fold) and quiescent regions (3.6-fold), and 8.7-fold depletion near TSSs (Supplementary file 1). We hypothesize that mutations affecting accessibility near TSSs are more likely to be deleterious, and thus selected against, resulting in the observed depletion. This 8.7-fold depletion near TSSs is greater than the analogous depletion of nonsynonymous changes in exons; in fact, we estimated that that these caQTLs are ~81% more likely to be deleterious, and removed by selection, than nonsynonymous mutations (see Supplemental Note).

**Figure 3.**
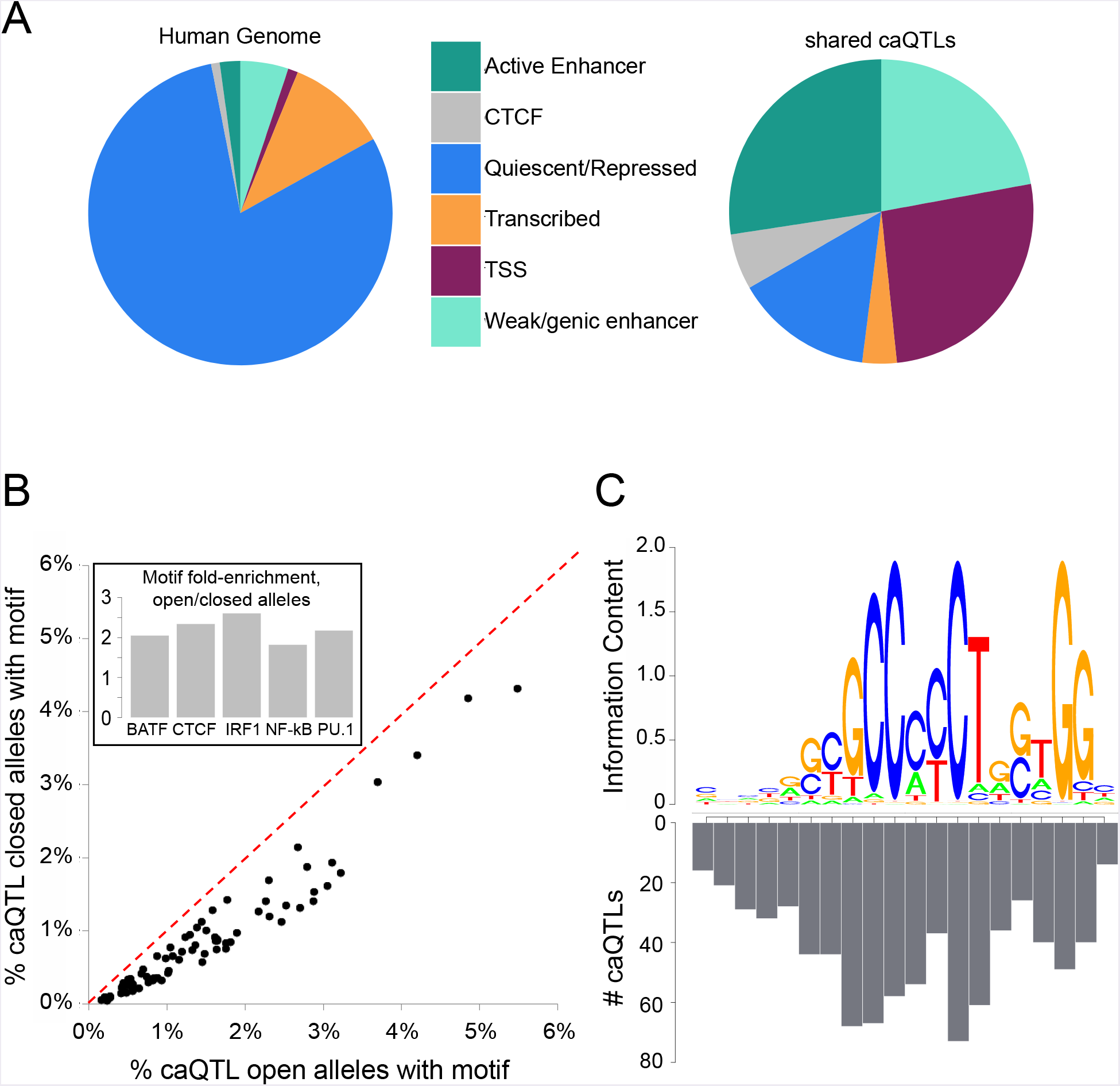
Characterizing shared caQTLs. A. The fraction of the genome (left) and of shared caQTLs (right) in each of four classes, annotated based on chromatin signatures (Ernst and Kellis 2012). TSS includes TSS flanking regions; full results in Supplementary file 3. **B**. Searching for motifs enriched specifically among open alleles (using closed alleles from the same caQTLs as the background comparison set), we found 80 motifs enriched among open alleles (points below the diagonal). Repeating the analysis for closed alleles, we found no motifs enriched (above diagonal). Note that many motifs are partially overlapping, and thus not independent. Inset: fold-enrichment in open/closed alleles for five selected TFs. Full results in Supplementary file 1. **C**. The number of caQTLs overlapping each position within the CTCF motif strongly mirrors the information content (i.e. the importance for binding) of that position, as expected if these caQTLs are causal variants affecting CA via CTCF binding. Full results in Supplementary file 1.

Disruption of TF binding motifs can lead to caQTLs (Degner et al. 2012, Kumasaka et al 2016). To investigate the effects of caQTL variants on TF binding motifs we searched for motifs enriched specifically in the more accessible caQTL alleles (by using the less accessible alleles as the background set), or in the less accessible alleles (see Methods). We found 80 known TF motifs enriched in the open allele sequences (FDR ≤ 0.1% for each; Figure 3B; Supplementary file 1), but none enriched in the closed alleles from the same caQTL loci. This striking asymmetry supports the idea that a major mechanism leading to caQTLs is disruption of TF binding, where caQTL variants matching the consensus motif—and thus promoting TF binding—result in more accessible chromatin. The most highly enriched motifs included TFs specific to immune cells like BATF, as well as more ubiquitous factors like CTCF ( Figure 3B inset).

To further investigate fine-mapped caQTLs in TF binding motifs, we focused on CTCF, due to its long and well-characterized motif (Ding et al. 2012). We reasoned that the most critical positions in the CTCF motif (i.e. those with the least variation across CTCF binding sites, represented by high information content in the position weight matrix) should be more likely to affect CTCF binding and chromatin when disrupted, and thus should be the most enriched for caQTLs (Ding et al. 2012, Maurano et al. 2015). Comparing shared caQTL density with the CTCF motif, we indeed found that the caQTL density is strongly correlated with the information content of each motif position (Pearson r=0.86; Figure 3C).

In sum, our analyses of the fine-mapped caQTLs support the idea that these are highly enriched for causal variants, since non-causal SNPs in LD with causal variants would not be expected to disrupt known motifs ( Figure 3B) or critical motif positions ( Figure 3C). Our results reveal both simple (trinucleotide) and complex (TF binding site) context-dependencies of variants affecting CA.

### Chromatin accessibility impacts long-range chromosomal interactions

CTCF has a well-established role in mediating long-range interactions between chromosomal loci, an essential component of transcriptional regulation (Guo et al. 2015, Sanborn et al. 2015, Rao et al. 2014). Consistent with this role, we previously reported that bQTL alleles increasing CTCF binding also increase these long-range interactions (Tehranchi et al. 2016). To test if caQTLs also affect long-range interactions, we measured how often the more accessible allele had significantly more long-range interactions (with loci >20 kb away) (Rao et al. 2014) than the less accessible allele, and vice versa; we found a 2.5-fold enrichment of the more accessible allele having more interactions ( Figure 4A; binomial p < 10^-36^), similar to the 2.2-fold bias of CTCF bQTLs (Tehranchi et al. 2016). The allelic ratio increased slightly (to 2.7-fold) when restricted to interactions >100 kb apart. This enrichment was not observed for inter-chromosomal interactions (Figure 4--figure supplement 1), suggesting it is unlikely to be due to a nonspecific bias in the Hi-C assay. These results establish a role for CA in polymorphic long-range chromosomal interactions of a similar magnitude as CTCF.

**Figure 4.**
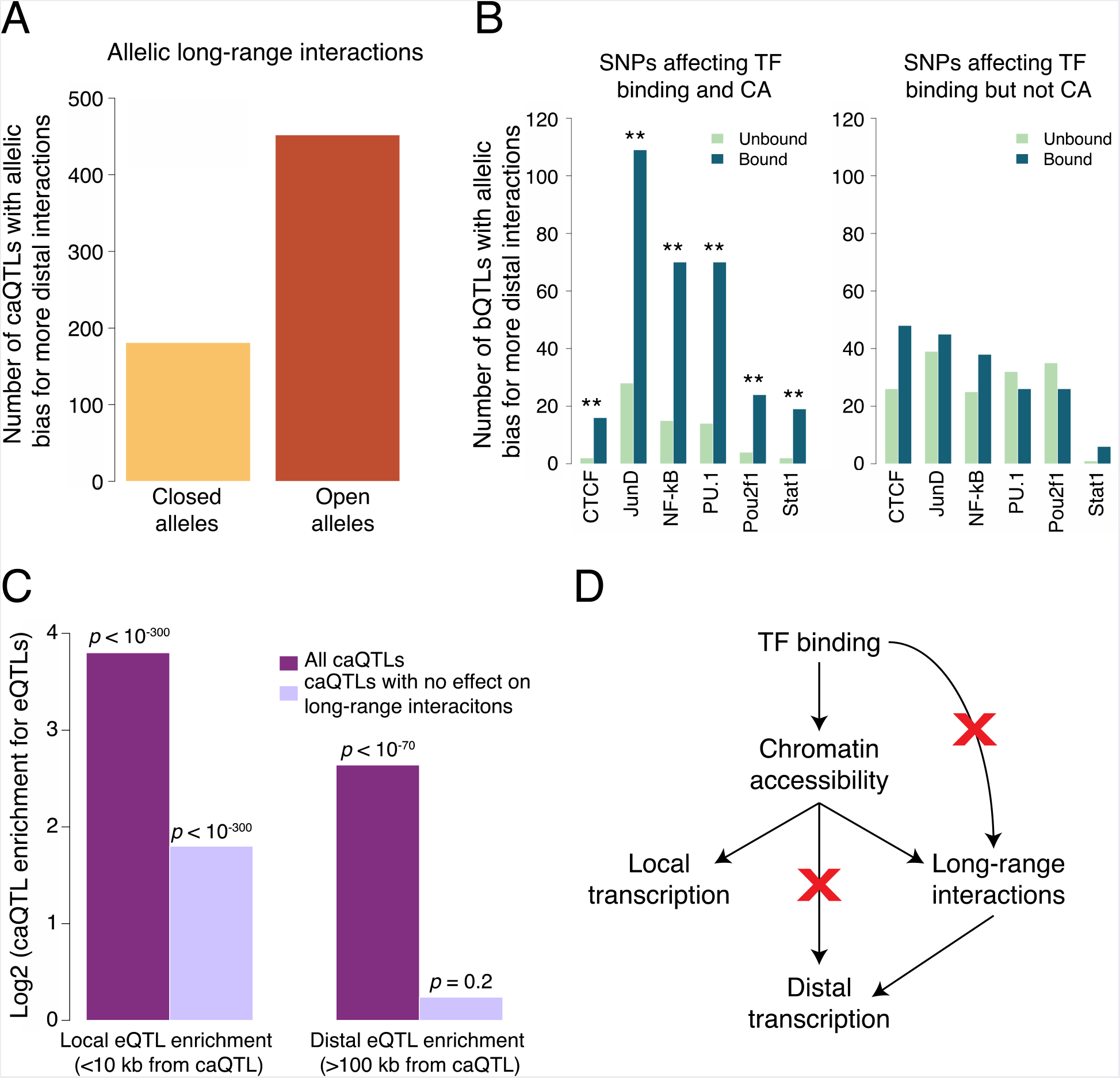
TF binding and chromatin accessibility. A. Using allele-specific 3D chromosomal interaction (Hi-C) data from an LCL (Rao et al. 2014), we found that open alleles of caQTLs tend to have more long-range interactions than do the closed alleles, establishing a role for CA in polymorphic chromosomal interactions. **B**. Splitting bQTLs into two groups (Figure 4--figure supplement 2), we found that bQTLs were strongly associated with the extent of long-range interactions only when they also affect CA (left panel; ** indicates Bonferroni-corrected binomial p < 0.008 for all six TFs); for bQTLs that do not affect CA, no allelic bias was observed (right panel; Bonferroni-corrected binomial p > 0.08 for all six TFs). **C**. caQTLs are strongly enriched for both local and distal eQTLs; however among those that do not affect long-range chromosomal interactions, only local eQTLs are enriched. **D**. Model summary: our results suggest that bQTLs generally cannot affect long-range chromosomal interactions without an effect on CA, and caQTLs generally cannot affect distal transcription without an effect on long-range interactions. The model shown represents a plausible interpretation, but is not the only possible causal scenario.

Our previous work showed that in addition to CTCF bQTLs, bQTLs for five other TFs also affect long-range chromosomal interactions, suggesting a possible role for many TFs in chromosomal architecture (Tehranchi et al. 2016); however, the underlying mechanism and causality has not been investigated. We hypothesized that we could isolate the effects of TF binding and CA by separating bQTLs into two groups: those that affect TF binding and CA, and those that affect TF binding but not CA (Figure 4--figure supplement 2). Comparing the effects of these two groups on long-range interactions could then shed light on whether TFs can shape chromosomal interactions independent of CA, or if instead CA effects are essential for TFs to impact these interactions.

We found a strong and consistent effect on long-range interactions for bQTLs that affect accessibility, in which the allele with increased TF binding is biased towards having more interactions ( Figure 4B left; binomial p < 0.008 for each). In contrast, we found no significant effect for the bQTLs that do not affect accessibility, for all six TFs ( Figure 4B right; binomial p > 0.08 for each). This suggests that TF binding alone has no detectable association with long-range interactions; even for CTCF, an effect on CA is also necessary. Together with additional evidence (See Supplemental Note), we propose that CA is a key intermediary between TF binding and chromosomal interactions ( Figure 4D).

### caQTLs that regulate distal genes alter the landscape of long-range interactions

We then asked whether these polymorphic chromosomal interactions are involved in the regulation of transcription by distal caQTLs (defined as a caQTL that is also an eQTL for a gene whose TSS is >100 kb away). Enhancers can act through physical interactions with promoters, and genetic variants can impact chromatin and transcription at distal loci with which they physically interact (Waszak et al. 2015, Grubert et al. 2015, Tehranchi et al. 2016). However, whether these long-range effects of SNPs depend on changes in the patterns of chromosomal interactions—as opposed to being mediated by static interactions—has not been investigated.

To address this question, we controlled for the effect of changes in long-range interactions by analyzing caQTLs with no measurable effect on these interactions (see Methods). If caQTL effects on distal genes do not depend on changes in these interactions, then eQTL enrichment should not be affected by controlling for them; if instead changes in these interactions are necessary for long-range regulation, then we would expect to see a sharp reduction in eQTL enrichment when controlling for them.

As expected, we found that overall caQTLs were strongly enriched for local eQTLs (TSS <10 kb from the caQTL); restricting the analysis to caQTLs with no effect on interactions reduced the magnitude of this enrichment, but it was still highly significant in both cases ( Figure 4C left; Fisher’s exact p < 10’^3^°° for each enrichment), suggesting that changes in long-range interactions are not necessary for caQTLs to affect transcription of nearby genes. caQTLs were also enriched for distal eQTLs (p < 10’^70^), but this enrichment was entirely lost when restricting the analysis to caQTLs with no change in long-range interactions ( Figure 4C right; p = 0.20). This suggests that alterations in the patterns of long-range chromosomal interactions are necessary for caQTLs to affect distal gene expression ( Figure 4D), and therefore that these polymorphic interactions may be an important mechanism by which eQTLs can act over vast genomic distances.

### Fine-mapping GWAS associations with caQTLs

In addition to revealing insights into transcriptional regulation, caQTLs also provide a means to explore genotype/phenotype associations by identifying likely causal variants and their molecular mechanisms of action. We illustrate different types of insights with three examples from autoimmune diseases, the class of disease most directly relevant to LCLs. In rare cases, a lack of LD allows GWAS to implicate a single likely causal variant; this is the case for rs4409785, which is associated with rheumatoid arthritis, multiple sclerosis, and vitiligo (International Multiple Sclerosis Genetics Consortium 2011, Jin et al. 2012, Okada et al. 2014), and was predicted to be the causal variant for vitiligo with 85% probability (Farh et al. 2015). The SNP is bound by CTCF in LCLs (ENCODE Project Consortium 2012) and disrupts a CTCF motif; we found that it is a caQTL in two European populations (FIN and TSI, p < 3x10^-6^ in each; not tested in African or ASW populations due to its low MAF), thus providing evidence of an effect on accessibility that is likely mediated by CTCF.

In other cases, multiple SNPs in LD have conflicting evidence of causality. For example, rs479844 is a lead SNP associated with atopic dermatitis in Europeans, and although it was assigned a 90% probability of being causal (Farh et al. 2015), it has not replicated in some populations (Lepre et al. 2013). A meta-analysis reported that when non-European ethnicities are included, the only consistent association is for another nearby SNP, rs10791824 (Paternoster et al. 2015). This variant is a strong caQTL in all six non-European populations (p < 10^-42^ in each) and also significant in 2/4 European populations (p < 5x10^-4^ in each), supporting the trans-ethnic GWAS (Paternoster et al. 2015) implicating it as the likely causal variant. In addition, the caQTL is an eQTL for *OVOL1* (Lappalainen et al. 2013, GTEx Consortium 2017), a TF involved in skin development, and has been found to affect transcription in a large-scale reporter assay in LCLs (Tewhey et al. 2016).

The third and most frequent case is where many SNPs are in strong LD, and thus have almost equally strong associations with a disease. For example, a large LD block of 62 variants on chromosome 3 is associated with multiple sclerosis, all of which have <5% probability of being causal based on GWAS signal alone (Farh et al. 2015). However one of these, rs485789, is a caQTL shared across 8/10 populations (p < 2x10^-5^ in each). Interestingly, this was also predicted as a causal variant by a sequence-based predictor of regulatory variants (Lee et al. 2015), and is an eQTL for *IL12A* (Lappalainen et al. 2013), a cytokine implicated in several autoimmune diseases (Guo et al. 2016). Therefore, rs485789 is a likely causal variant that acts on *IL12A* via its effect on CA.

More broadly, we found 5598 caQTLs that were also associated with disease risk or other complex traits (GWAS p < 5x10^-8^; Supplementary file 4). Although most GWAS loci include dozens of potential causal variants in LD, there are only ~2.2 caQTLs per GWAS locus, providing a far smaller credible set for targeted follow-up studies. Among these, 115 caQTLs were shared across all ten populations, suggesting that many of these are likely to be causal for disease risk; we highlight ten examples in Table 1. We note that although our caQTLs were measured in LCLs, the traits they are associated with are related to a wide range of tissues. Consistent with this, we found that the regulatory effects of caQTLs are typically shared across most tissues, suggesting that their effects on CA are broadly shared as well (Supplemental Note, Table 1--table supplement 1). Therefore, it should not be surprising that these caQTLs can contribute to risk for diseases that have no clear connection to LCLs.

**Table 1.** Ten candidate causal variants, shared as caQTLs across all 10 populations. GWAS information is from the GRASP database (Eicher et al. 2015). See Supplementary file 4 for all caQTL/GWAS overlaps.

| <u>chr</u> | <u>caQTL position</u> | <u>caQTL rsID</u> | <u>GWAS rsID</u> | <u>GWAS p-value</u> | <u>GWAS trait</u> | <u>Candidate gene</u> |
| --- | --- | --- | --- | --- | --- | --- |
| 15 | 86018746 | rs7161880 | rs4281668 | 9.90E-09 | Aggressive prostate cancer | *AKAP13* |
| 9 | 5097544 | rs3780372 | rs10974944 | 5.10E-32 | Myeloproliferative neoplasms | *JAK2* |
| 12 | 111351439 | rs11065769 | rs10849917 | 3.21E-08 | Coronary artery disease | *MYL2* |
| 2 | 68598955 | rs17035378 | rs17035378 | 8.00E-09 | Celiac disease | *PLEK* |
| 6 | 167433948 | rs6904946 | rs2301436 | 1.00E-12 | Crohn's disease | *RNASET2* |
| 14 | 69273905 | rs194749 | rs194749 | 2.70E-10 | Inflammatory bowel disease | *ZFP36L1* |
| 1 | 154359411 | rs9651053 | rs11265608 | 2.75E-08 | Juvenile idiopathic arthritis | *IL6R* |
| 4 | 1731653 | rs798764 | rs798766 | 3.90E-13 | Urinary bladder cancer | *FGFR3* |
| 13 | 40334852 | rs9603612 | rs9532434 | 4.52E-08 | Juvenile idiopathic arthritis | *COG6* |
| 11 | 118560857 | rs73001406 | rs11216930 | 1.40E-09 | Glioma | *PHLDB1* |

## Discussion

By applying pooled QTL mapping in 1000 cell lines from 10 diverse populations, we achieved unprecedented power and resolution to fine-map cis-regulatory variants. The resulting collection of over 100,000 caQTLs allowed us to refine our understanding of the sequence-level causes, as well as the phenotypic consequences, of natural variation in chromatin accessibility.

More generally, our results show that molecular QTL mapping and trans-ethnic fine-mapping are complementary in several respects. Molecular QTL mapping provides information about molecular mechanisms and allelic effect direction, while including multiple populations allows for fine-mapping of causal variants; combined, these can provide unprecedented insights into the genetic basis of human variation. Moreover, since variation in molecular-level traits such as CA can be mapped to over 10^5^ QTLs, they provide tremendous opportunities for fine-mapping regulatory variants throughout the genome, which can then inform follow-up studies for diseases or other traits where trans-ethnic fine-mapping may not be feasible (e.g. Table 1). We expect that future studies will greatly benefit from this combination, and that the ease of pooled QTL mapping will facilitate the adoption of this integrated approach.

As a first step towards understanding how caQTLs impact phenotypes, we have explored their effects on local and distal transcriptional regulation. This strategy is conceptually similar to experiments involving genome editing of specific regulatory elements (Guo et al. 2015, Sanborn et al. 2015), except that we are utilizing natural genetic variation as our source of perturbations; this allows us to examine many more variants, though we are restricted to those that are not strongly deleterious (since these are too rare for us to observe). Our results suggest that TF binding affects long-range chromosomal interactions via its effect on CA, and that CA affects transcription of distal genes via its effect on long-range chromosomal interactions (Figure 4). This latter inference is especially intriguing, since it suggests a molecular mechanism underlying distal eQTLs: variants alter enhancer activities, which in turn alter the physical interactions of enhancers with distal genes. To our knowledge, this represents the first evidence that variation in the landscape of chromosomal interactions underlies long-range genetic effects on transcription.

There are several important limitations to our study. First, we have yet to map the majority of human caQTL variants—e.g. those specific to other cell types, those that affect CA of distal loci (in *trans* or long-range cis), or those involving rare variants or small effect sizes. Second, our shared caQTLs should be regarded only as candidate causal variants to guide future experiments, similar to other statistically fine-mapped associations. And finally, much work remains to understand how the caQTLs we have mapped affect human phenotypes.

Looking ahead, we expect that combining GWAS with pooled QTLs mapped in diverse populations and cell types will be an efficient and highly effective strategy for identifying causal variants and their molecular mechanisms of action, two major bottlenecks in bridging the gap between GWAS and disease etiologies.

## Materials and Methods

### Cell culture conditions

Lymphoblastoid cell lines (LCLs) from unrelated individuals were obtained from the Coriell Institute (HYPERLINK "http://www.coriell.org"). The LCLs were grown in RPMI-GlutaMAX-HEPES media (Life Technologies, 72400) with 15% FBS, 100 I.U./mL penicillin, and 100 μg/Ml streptomycin at 37°C, 5% CO_2_.

### ATAC-seq

We used a modified version of the ATAC-seq protocol (Buenrostro et al. 2015). Each individual cell line was grown to a density of 6-8x10^5^ cells/mL and 2x10^3^ cells were collected and pooled by population such that each individual was approximately equally represented in the pool. ! Subpools were frozen in liquid nitrogen at -180°C. After all 1000 individuals were collected, subpools were combined by population and fresh media was added up to 15ml, centrifuged at 1500rpm for 10min, and supernatant removed. The pellet was resuspended in 0.75ml media with 200 units/mL DNase and incubated in a ThermoMixer for 30 minutes at 37°C at 300rpm. 0.8mL of Ficoll-paque Plus (GE Cat #17-1440-03) was added to a 2mL tube and the 0.75mL cel suspension was carefully layered on top. Samples were centrifuged at 500*x g* for 20 min at room ! temperature with no brake. The thin cloud of live cells in the middle layer were pipetted to a new tube, washed in 1ml 1x PBS buffer, and split into two tubes with 10^5^ cells for replicates. Tubes Iwere centrifuged at 500×g for 5 min at 4°C and supernatant was removed.

ATAC-seq was performed simultaneously on all 20 replicates. To each cell pellet, 100ul of ! transposition mix was added (50uL 2x TD Buffer (Illumina Cat #FC-121-1030), 5.0 uL Tn5 Transposase (Illumina Cat #FC-121-1030), 42uL nuclease-free water, 1uL 10% Tween-20, 3uL 1% Digitonin). Samples were incubated in a ThermoMixer at 37°C for 30 min@750rpm, then purified using Qiagen MinElute Kit with DNA eluted in 11 uL 10mM Tris buffer, pH 8. Transposed DNA fragments were amplified using PCR where total cycles were calculated using qPCR as described in (Buenrostro et al. 2015). Amplified libraries were purified using Qiagen PCR Cleanup Kit and eluted in 21uL 10mM Tris pH 8. An additional purification step was performed using a 1:1.2 ratio of DNA:AMPure XP beads. Libraries were sequenced on an Illumina HiSeq 4000 (150 bp, paired-end reads).

### Mapping ATAC-seq reads

To remove adapters, reads were trimmed using cutadapt (Martin 2011) with the following command: cutadapt -e 0.20 -a CTGTCTCTTATACACATCT -A CTGTCTCTTATACACATCT -m 5 -o fastq1out -p fastq2out fastq1 fastq2

Trimmed reads were mapped using a modified version of the WASP pipeline for controlling mapping bias (van de Geijn et al. 2015) with scripts find_intersecting_snps_2.py and filter_remapped_reads_2.py that can be found at:HYPERLINK https://github.com/TheFraserLab/WASP/tree/atac-seq-analysis/mapping". Briefly, for each read overlapping a SNP, we remapped hypothetical reads with the other allele, and discarded any reads that do not map uniquely, to the same location, for both alleles. Duplicate reads were filtered out for each replicate using HYPERLINK "https://github.com/eilon-s/bioinfo_scripts/rmdup.py".

### Mapping and analyzing caQTLs

Pre- and post-ATAC allele frequencies, and the resulting p-values, were calculated using our published pipeline (Tehranchi et al. 2016). This uses post-ATAC allele frequencies together with individual sample genotypes to infer pre-ATAC allele frequencies.

To estimate pre-ATAC allele frequencies, the pre-ATAC pool could be sequenced; however this suffers from two major drawbacks. First, although sequencing the pre-ATAC pool could yield accurate pre-ATAC allele frequencies, it cannot account for genome-wide differences between samples such as the total amount of open chromatin. If one sample has more open chromatin than another, its alleles will be over-represented in the post-ATAC fraction. This will constitute a source of noise in our analysis, since our goal is to map cis-acting variation affecting individual sites. Second, sequencing the pre-ATAC pool would require very deep sequencing to achieve accurate allele frequency estimates at each potential caQTL SNP, since the sequencing would not be restricted to SNPs in open chromatin regions (as they are for the ATAC fraction).

Therefore, we previously developed an alternative which does not require any additional sequencing, and does account for genome-wide differences between samples. Our regression-based approach (Tehranchi et al. 2016) uses the post-ATAC frequencies together with genotypes of each sample to infer the proportion of each sample in the pool. These proportions will be weighted by any genome-wide differences, since these will be naturally incorporated into the post-ATAC frequencies used as input to the regression. In this way, our pre-ATAC allele frequencies already account for some types of *trans*-acting variation, increasing our power for mapping *cis*-acting differences.

SNPs were considered testable in a population if they were covered by at least 20 ATAC-seq reads and had minor allele frequency > 0.01 in that population. No peak calling was performed; each SNP was tested for allelic bias using only the reads overlapping it, since these reads are the only ones that give information about its allelic bias (Tehranchi et al. 2016). Genotypes were downloaded from the 1000 Genomes Project. Scripts and documentation can be found at: HYPERLINK "https://github.com/tehranchi/public".

Directionality and enrichment tests were performed as described in (Tehranchi et al. 2016). Numbers underlying specific analyses are reported in Supplementary file 1.

### Fisher’s combined probability test

Shared caQTLs were calculated using Fisher’s combined probability test, where*p_t_* is the caQTL p-value for population i. Any SNP not tested in population *i* was assigned*p_i_* = 1.

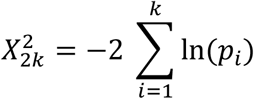

Nearly all (99.8%) of the shared caQTLs at Fisher’s combined p<5x10^-6^ were at least nominally (p<0.05) significant across multiple populations, and 98.6% showed concordance of the more accessible allele between the top two populations.

### Motif analysis

We used HOMER (Heinz et al. 2010) to search for motifs that were differentially enriched among the high-CA alleles compared to the low-CA alleles in our shared caQTLs. The caQTL plus 15 bp on each side (31 bp total) was used as input to HOMER, with the less accessible alleles of the same caQTLs used as the background comparison set. Therefore all significant enrichments are due to caQTL variants within the motifs themselves (since any motifs flanking the caQTLs would be present in both comparison sets). We ran the findMotifs.pl script from the HOMER package where the targetSequences.fa file contained the high-CA alleles and the background.fa file contained the low-CA alleles, and then repeated the analysis with the two files swapped: findMotifs.pl <targetSequences.fa> fasta <output directory> -fasta <background.fa>

### Long-range interaction analysis

To obtain Fig. 4A-B, we restricted the analysis to shared caQTLs (Fisher’s combined p < 5x10^-6^) with consistent directionality in a majority of the populations tested, and counted the number of Hi-C reads connecting each caQTL allele with any distal locus (>20 kb or >100 kb away). We then determined the number of caQTLs where the more accessible allele had significantly more Hi-C contacts (using a binomial test of allele-specific read counts), compared to the number where the less accessible allele was more interactive. See Supplementary file 1 for detailed results.

For Fig. 4C we first obtained two subsets of eQTLs (Lappalainen et al. 2013): those that are proximal to their target TSSs (< 10 kb) and those that are distal (> 100 kb). Next, we calculated the enrichments of these proximal and distal subsets among shared caQTLs. To obtain the p-values in Fig. 4C, we used Fisher’s exact test to compare the significance of these enrichments with the respective enrichments of proximal and distal eQTLs among non-caQTLs (Fisher’s combined p > 0.5). We repeated this analysis on the subsets of caQTLs and non-caQTLs in regions without significant allelic bias (binomial p > 0.5) in the number of reads supporting long range (> 100 kb) interactions, requiring each SNP to have at least 100 reads supporting long-range interactions in order to be confident in the lack of strong allelic bias. The non-significant (p = 0.20) enrichment in Fig. 4C is based on a relatively large number of SNPs (1680 caQTLs with no effect on interactions, and 5719 non-caQTLs with no effect on interactions)—the same background set as used for the significant (p < 10^-70^) enrichment in the same figure—suggesting that a small sample size is not driving the lack of significant enrichment. See Supplementary file 1 for further details of this analysis.

### GWAS overlaps

We used all reported GWAS associations at p < 5x10^-8^ from the GRASP database (28), excluding those for gene expression levels (eQTLs). In order to include cases where the reported SNP is not the causal variant, we expanded the GWAS list to include any variants in strong LD with GWAS hits (r^2^ > 0.8 in the CEU population; CEU is similar to the European cohorts in which nearly all of the GWAS were conducted). We then intersected this LD-expanded list with our caQTLs (Supplementary file 4). Because most GWAS hits are in strong LD with other variants, many of the caQTLs that overlap GWAS hits may not be causal SNPs for the GWAS trait; we cannot apply published co-localization methods to quantify this, since these require association data across an entire locus or LD block for comparison, and our pooled approach is limited to SNPs covered by the ATAC-seq reads.

### IDR analysis

For IDR calculation, we used parameters similar to the ENCODE project (mu=0.1, sigma=1.0, rho=0.2, p=0.5, eps=10^-6^, max.ite=3000). Since IDR does not allow tied p-values, we broke ties by adding a small random number (normally distributed with standard deviation 10^-17^).

## Supplementary Text

### Assessing caQTL reproducibility with IDR

A standard approach for estimating an FDR in association studies is to randomize the data, e.g. randomly pairing one sample’s genotypes with another sample’s ATAC-seq data, and then recalculate the associations to gauge the extent of false positives. Since our approach does not have separate data for each individual, such randomization is not possible, and thus we cannot estimate an FDR for our caQTLs using standard approaches. However, we can apply a related method known as the irreproducible discovery rate (IDR), developed as part of the ENCODE project (Li et al. 2011, ENCODE Project Consortium 2012) to assess the statistical reproducibility of our caQTL p-values. To measure this, we calculated caQTL p-values separately for each biological replicate of two populations, ASW and CEU. The IDR estimates the fraction of data points that are not reproducible at any given p-value cutoff. For a range of potential cutoffs, we observed a similar trend in both populations: at p = 5x10^-3^ we observed IDR ≈ 0.03, and p = 5x10^-4^ corresponds to IDR ≈ 0.01 (dashed red line in Figure 1—figure supplement 3). These numbers are consistent with other results suggesting a low FDR among our caQTLs, such as their ~97-99% agreement with dsQTL and bQTL directionality (Fig. 1E-F).

### Comparison to non-pooled chromatin accessibility studies

In our comparison with RASQUAL caQTLs (Kumasaka et al. 2016), we first sought to identify a matched significance cutoff. Among their 2,707 caQTLs (FDR=10%), 3.5% were previously identified as dsQTLs (Degner et al. 2012) (the most closely matched type of QTL available for comparison). This 3.5% overlap between two previous studies of chromatin accessibility QTLs may seem low, but likely results from differences in experimental methods (ATAC-seq vs DNase-seq), analysis methods, and populations, as well as incomplete power. In CEU, our population most closely matched to the British (GBR) population used for RASQUAL, we found that our top 7587 caQTLs (p<3x10^-6^) yielded the same overlap of 3.5% dsQTLs. Our approach does not involve calling CA peaks, but in a separate analysis we estimated that <20% of our caQTLs fall into the same peak as another caQTL, so removing these would not have a major effect on the results.

In addition to the RASQUAL comparison, we also compared our caQTLs to variants associated with allelic imbalance of chromatin accessibility (Maurano et al. 2015). In that study, 493 samples from 166 individuals were profiled with a total of 2.6x10^10^ DNase-seq reads. At the most stringent cutoff (FDR=0), they called 2,420 imbalanced SNPs, 2.1% of which were also dsQTLs. Comparing this to our study, our top 16,990 CEU caQTLs (p<3.5x10^-4^) had the same 2.1% enrichment. Therefore we mapped 7-fold more caQTLs from 246-fold fewer libraries and 443-fold fewer reads, though their inclusion of diverse tissues likely decreased the overlap with dsQTLs from LCLs.

We note that dsQTLs are not the only possible benchmark we could use to select caQTL lists of equal quality across studies. When using CTCF bQTLs (Ding et al. 2014) instead, our estimated improvement in cost per QTL was even greater than when using dsQTLs, due to the high enrichment of CTCF bQTLs in our data (~190-fold in our CEU caQTLs [Figure 1—figure supplement 5], compared to ~7-fold in RASQUAL caQTLs [Kumasaka et al. 2016] and 4.5-fold in the imbalanced DNase-seq SNPs [Maurano et al. 2015]).

### Comparing selection against caQTLs and nonsynonymous variants

If this depletion of caQTLs near TSSs (Fig. 3A) is due to selection against deleterious effects of these caQTLs, then we can compare this with human Ka/Ks ratios, which reflect the depletion of nonsynonymous changes in protein-coding regions. Across all human genes, Ka/Ks since divergence with chimpanzee was estimated as 0.208 (Chimpanzee Sequencing and Analysis Consortium 2005), implying ~4.8-fold depletion for nonsynonymous changes. Therefore we observed 8.7/4.8 = 1.81-fold (i.e. 81%) greater depletion for the caQTLs. This comparison is only an approximation, since it ignores effects of positive selection, as well as the possibility that caQTLs are depleted near TSSs for reasons other than selection (e.g. CA having greater robustness to mutations near TSSs). We also note that if there is any negative selection against TSS-proximal SNPs that are not caQTLs then the 1.81-fold difference will be an underestimate.

### Effects of caQTLs on DNA shape

Since the shape of the DNA double helix is sequence-dependent, and can affect interactions with TFs, we tested whether shared caQTL variants tend to cause larger changes in DNA shape than non-caQTLs. We defined a non-caQTL as a variant with caQTL p = 1 and at least 20 ATAC-seq reads. We first computed the total read depth for each shared caQTL and non-caQTL across all populations. We then created a read-matched set of shared caQTLs and non-caQTLs in the following way: For each shared caQTL, we identified a unique non-caQTL where | log_2_(shared caQTL reads) -log_2_(non-caQTL reads) | < 0.5. If no such non-caQTL existed, we eliminated the shared caQTL. We were able to find unique read-matched non-caQTLs for 11,416 of the shared caQTLs.

After identifying read-matched shared caQTLs and non-caQTLs, we used BEDTools version 2.26.0 (Quinlan and Hall 2010) to extract the genome sequence at each variant +/-5bp. For each shared caQTL, we created four sequences - one with the open allele +/-5bp, one with the closed allele +/-5bp, one with the read-matched non-caQTL reference allele +/-5bp, and one with the read-matched non-caQTL alternate allele +/-5bp. We then used DNAshapeR version 1.0.2 (Chiu et al. 2016) to estimate the minor groove width (MGW), propeller twist (ProT), Roll, and helix twist (HelT) for each of the sequences for each shared caQTL. We computed the difference in each shape parameter between the sequences for the open and closed caQTL alleles as well as between alleles for the read-matched non-caQTLs, where the “open” and “closed” alleles were randomly selected. To compute whether the shared caQTLs have a stronger association with DNA shape than the read-matched non-caQTLs, we used the Wilcoxon rank-sum test to compare the two difference distributions at the position of the variant. Since Roll and HelT shapes are computed in groups of two nucleotides, we concatenated the difference distributions for the two windows of 2 bp that overlap the variant. We multiplied each p-value by 4 as a Bonferroni correction.

The results show small but significant differences for three of the DNA shape measures (Figure 4—figure supplement 4).

### Causality of joint bQTLs/caQTLs affecting long-range chromosomal interactions

In addition to CTCF, we previously reported an association between long-range chromosomal interactions and binding of five additional TFs in LCLs (Tehranchi et al. 2016). However, in that work we did not explore the mechanism by which this occurs or the direction of causality: it could be that TF binding impacts long-range interactions, though other causal scenarios are also possible. For example, joint bQTLs/caQTLs could be cases where a variant first affects TF binding and then CA as a result; or the variant’s direct effect could be on CA, which then changes the local landscape of TF binding; or there may be independent effects on TF binding and CA that are not causally linked (either via two distinct causal variants in linkage disequilibrium, or one variant with independent effects on TF binding and CA). In general, the strong enrichment for overlap between caQTLs and bQTLs suggests that causal independence is rare at these joint QTLs, since under such a model one would not expect to see either strong enrichment for overlap, or the nearly perfect concordance in directionality that we observed (~98-99%). In most cases we cannot distinguish between the first two scenarios, i.e. whether the TF binding or CA effect “comes first”; however in cases where a joint bQTL/caQTL disrupts a known TF binding motif, then we can infer that its primary effect is likely to be on TF binding to DNA, and thus any other effects—on CA or other traits—are likely to be downstream consequences of this.

We used this logic to ask whether TF binding affects long-range chromosomal interactions, by restricting our test for long-range interactions only to those caQTLs that disrupt a known TF binding motif and are present within a DNase hypersensitive footprint indicative of TF binding (Degner et al. 2012, Maurano et al. 2015); these caQTLs are most likely caused by the SNP’s direct effect on TF binding having a subsequent effect on CA. For these caQTLs, we observed a statistically indistinguishable enrichment for the more accessible allele having more interactions (63/24 = 2.6-fold), indicating that disruptions of TF binding lead to changes in both CA and long-range chromosomal interactions. We therefore show TF binding upstream of CA in Fig. 4D, though the reverse causality could also occur in some cases.

We could not test the possibility that some other variants may have a primary effect on CA and secondary effects on TF binding, since we cannot determine which caQTLs are not bQTLs for any TF, given the limited amount of bQTL information available (Figure 4—figure supplement 2).

### Regulatory effects of caQTLs across tissues

To explore the effects of caQTLs on gene expression across tissues, we intersected our caQTLs with eQTLs mapped in a meta-analysis across 44 tissues/cell types (GTEx Consortium 2017). Each eQTL was assigned a posterior probability of being active in each tissue, and following the GTEx authors, we required p > 0.9 (i.e. 90% probability) to call an eQTL as likely active in a given tissue. Overall, the median number of tissues per eQTL was 19 out of 44. In contrast, among 2833 caQTLs that were also eQTLs, the median number of active tissues was 25/44 (Wilcoxon p = 4x10^-21^). This implies that the underlying caQTL effects are probably shared across most tissues as well, with 25/44 being a likely underestimate of the extent of tissue sharing, since caQTLs present in a tissue do not necessarily act as eQTLs (as we have seen for LCLs).

We then asked in what tissues/cell types were our caQTLs most likely to affect gene expression. To measure this, we calculated the ratio of [fraction of joint eQTL&caQTL with posterior p > 0.9 in tissue X] / [fraction of all eQTLs with posterior p > 0.9 in tissue X]; values greater than one indicate that the joint eQTL&caQTLs are more likely to affect expression in tissue X than are eQTLs overall (this accounts for differences in numbers of eQTLs per tissue, which is largely driven by sample sizes). All 44 tissues had values > 1.06, with two clear outliers, LCLs and whole blood, having the highest chance of caQTLs affecting expression (Table 1—table supplement 1); this is not surprising because LCLs were used in our study and whole blood is the most similar tissue to LCLs in this dataset. At the other extreme, the ten sampled brain regions and testes had the 11 lowest ratios, indicating that caQTLs active in LCLs are slightly less likely to affect gene expression in these tissues.

### caQTLs in one cell type can underlie eQTLs specific to other cell types/conditions

Our multi-tissue analysis above (Table 1—table supplement 1) suggested that caQTLs from LCLs can be informative about transcriptional regulation in other cell types. We hypothesized that even in cases where a caQTL is not an eQTL in LCLs, if the effect on CA is preserved across other cell types or conditions, then a change in the expression of a trans-acting factor could cause a constitutive caQTL to be a condition-specific eQTL. Indeed, in our multi-tissue analysis, we observed 200 caQTLs that had a posterior p < 0.1 of being an eQTL in LCLs, but nevertheless were called as likely eQTLs (p > 0.9) in an average of 11.4 other tissues. We have also come across several examples of this outside of the GTEx data, and highlight two of these below.

A previous study identified a SNP, rs9806699, that acts as an eQTL for *GATM* in LCLs only after exposure to the statin drug simvastatin (Mangravite et al. 2013). This SNP was also associated with statin-induced myopathy in two cohorts, though the molecular mechanism underlying this condition-specific eQTL was not investigated. We found that this same SNP is also a caQTL in the GWD population (p = 6.3x10^-5^), which may underlie its statin-dependent transcriptional response.

Another example is a caQTL that is one of five linked variants associated with rheumatoid arthritis, all in strong LD (r^2^ > 0.95) and with similar GWAS p-values (1.3x10^-7^ < p < 3.1x10^-7^) that preclude fine-mapping (Farh et al. 2015). The caQTL is in a “super-enhancer” (long enhancers bound by many TFs) 6 kb upstream of *LBH*, a regulator of synoviocyte proliferation (Ekwall et al. 2015). This same variant was recently shown to causally affect *LBH* transcription in fibroblast-like synoviocytes (Hammaker et al. 2016), with the less accessible allele associated with reduced expression (as well as higher RA risk). This variant has not been found as an eQTL for *LBH* in LCLs (GTEx Consrtium 2017, Lappalainen et al. 2013).

These two examples illustrate how caQTLs mapped in one cell type/condition can have transcriptional effects that are specific to other cell types/conditions.

## Acknowledgements

We would like to thank members of the Fraser lab, J. Pritchard, and W. Greenleaf for helpful advice, and M. Simon for sharing lab space. This work was supported by NIH grant 2R01GM097171-05A1. All ATAC-seq reads are available at NCBI SRA, project ID PRJNA383900. All genome sequence data are available at HYPERLINK "http://www.internationalgenome.org.

## Author contributions

AKT performed experiments. HBF conceived the project and wrote the paper. All authors performed data analysis and edited the paper.

## Supplementary File

**Supplementary file 1**. Detailed results of analyses shown in the Figures. Each tab has a separate summary:

*caQTL summary:* Numbers of cell lines, reads, SNPs, and caQTLs

*Shared caQTL summary :* Number of populations in which each shared caQTL reached p<0.005

*Individual coefficients*: The coefficients inferred from our regression approach for inferring contribution of each sample to each pool

*Numbers of QTLs:* Total numbers of dsQTLs, bQTLs, and Hi-C GM12878 heterozygous SNPs used in our analyses

*Fig 1C-F:* Numbers going into Fig 1C-F

*Fig 2A:* All caQTLs testable (at least 20 reads and MAF>0.01) in all ten populations

*Fig 3A:* Numbers in Fig. 3A, and chromatin states for every caQTL

*Fig 3B:* Known TF binding motifs enriched in the more accessible caQTL alleles

*Fig 3C:* Numbers going into Fig 3C

*Fig 4A:* Numbers going into Fig 4A, including additional distance cutoffs

*Fig 4C:* Summary numbers going into Fig 4C, and for each caQTL also called as an eQTL, the distance between the caQTL and the target gene’s TSS

**Supplementary file 2**. All cell lines used, and all caQTLs at a nominal p < 0.005. The numbers of caQTLs are greater than those shown in Fig. 1B, since for Fig. 1B and the associated text we removed those in LD (*r*^2^ > 0.8 in YRI).

**Supplementary file 3**. Shared caQTLs at a Fisher’s combined p < 5x10^-6^.

**Supplementary file 4**. All caQTL/GWAS overlaps. Rows where the GWAS rsID is the same as the caQTL rsID indicate that the GWAS variant was itself a caQTL; rows where they do not match indicate the two variants are in LD (*r*^2^ > 0.8 in CEU). caQTLs with p < 0.005 were included.

**Figure 1—figure supplement 1.**
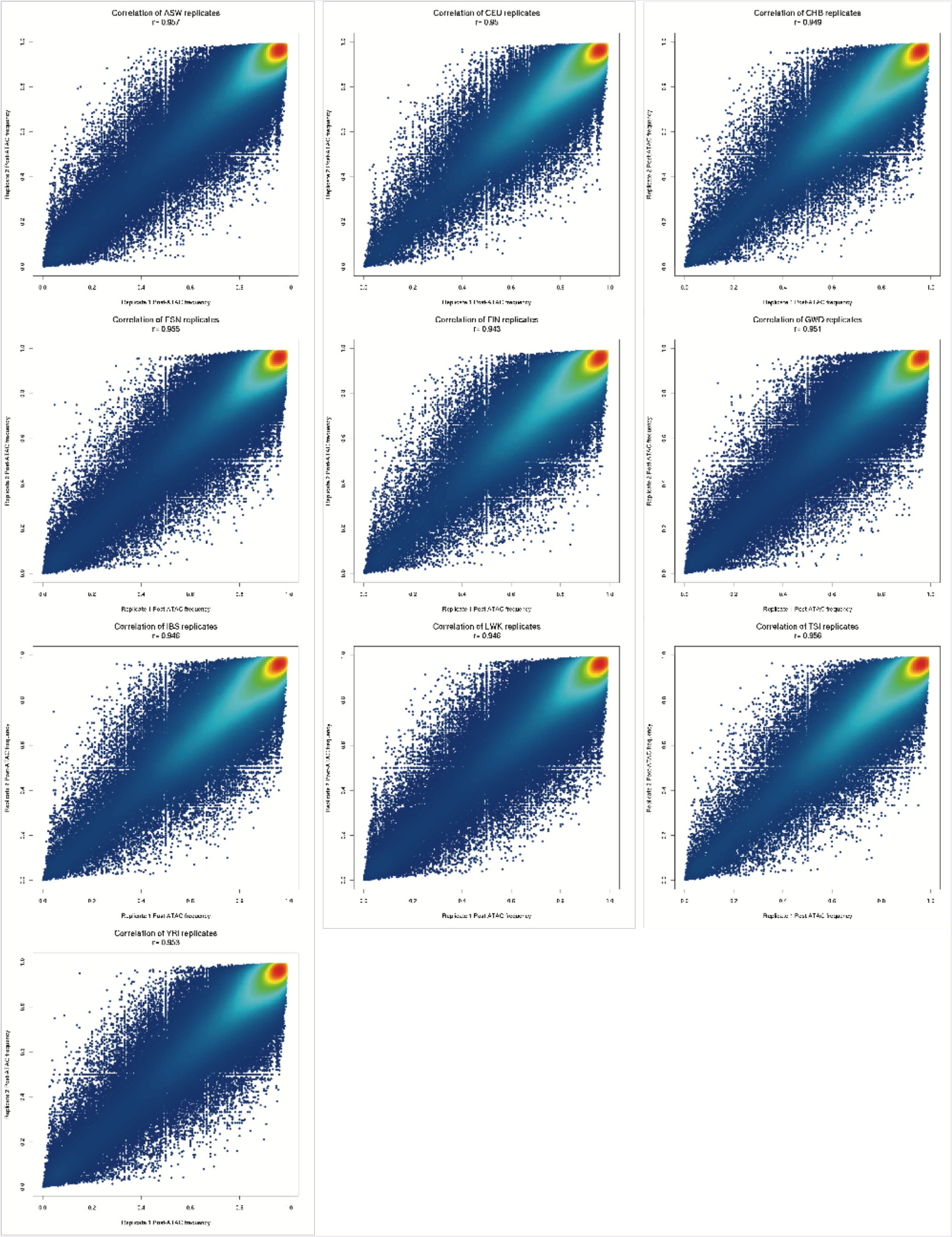
Comparison of post-ATAC reference allele frequencies between biological replicates of each population pool. All replicates have 0.94 < *r* < 0.96.

**Figure 1—figure supplement 2.**
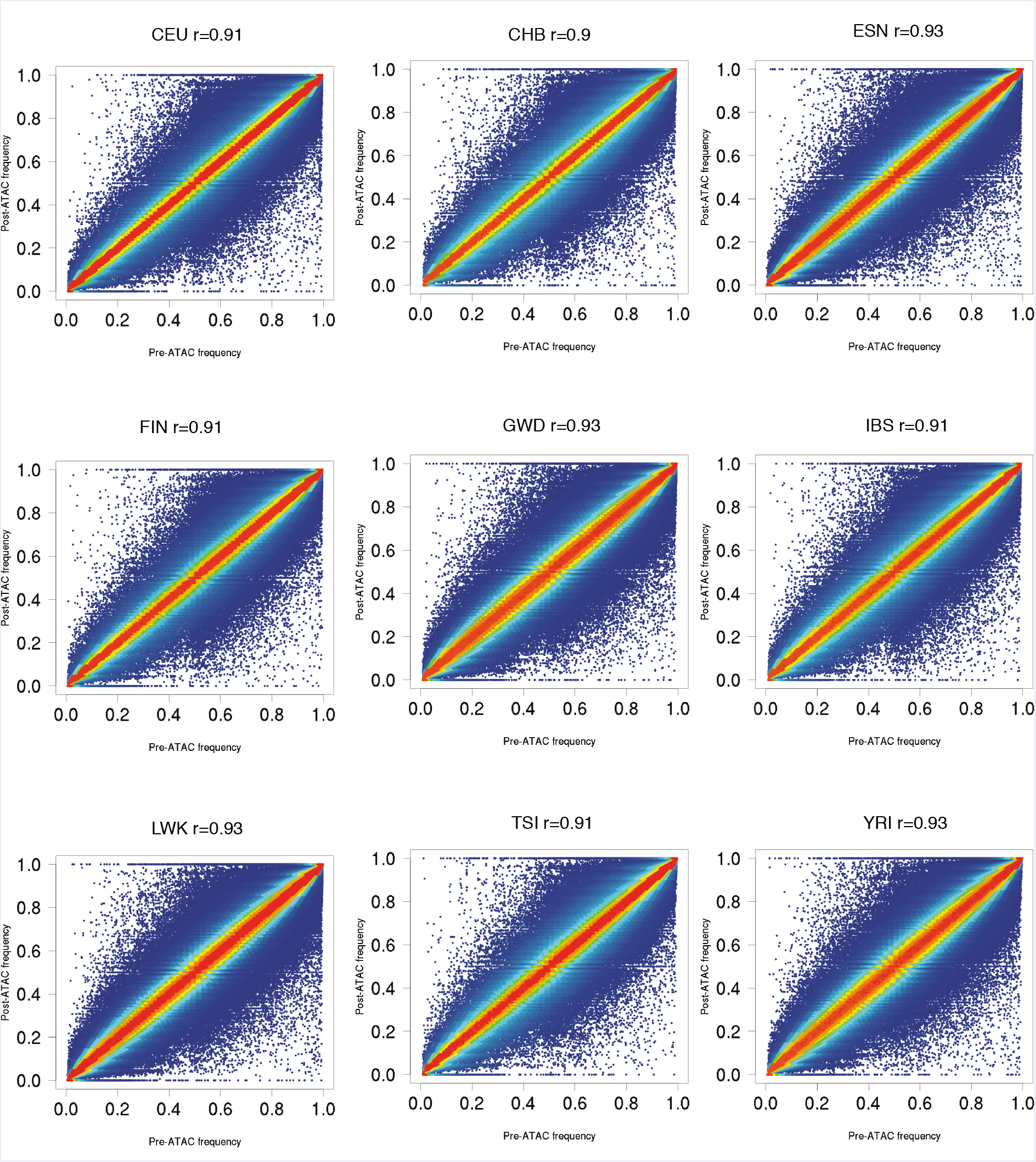
Pre-ATAC vs. post-ATAC reference allele frequencies for nine populations, similar to ASW plot in Fig. 1A. Most SNPs fall close to the diagonal, as expected if most SNPs are not caQTLs. All populations have 0.90 < *r* < 0.94.

**Figure 1—figure supplement 3. IDR analysis.**
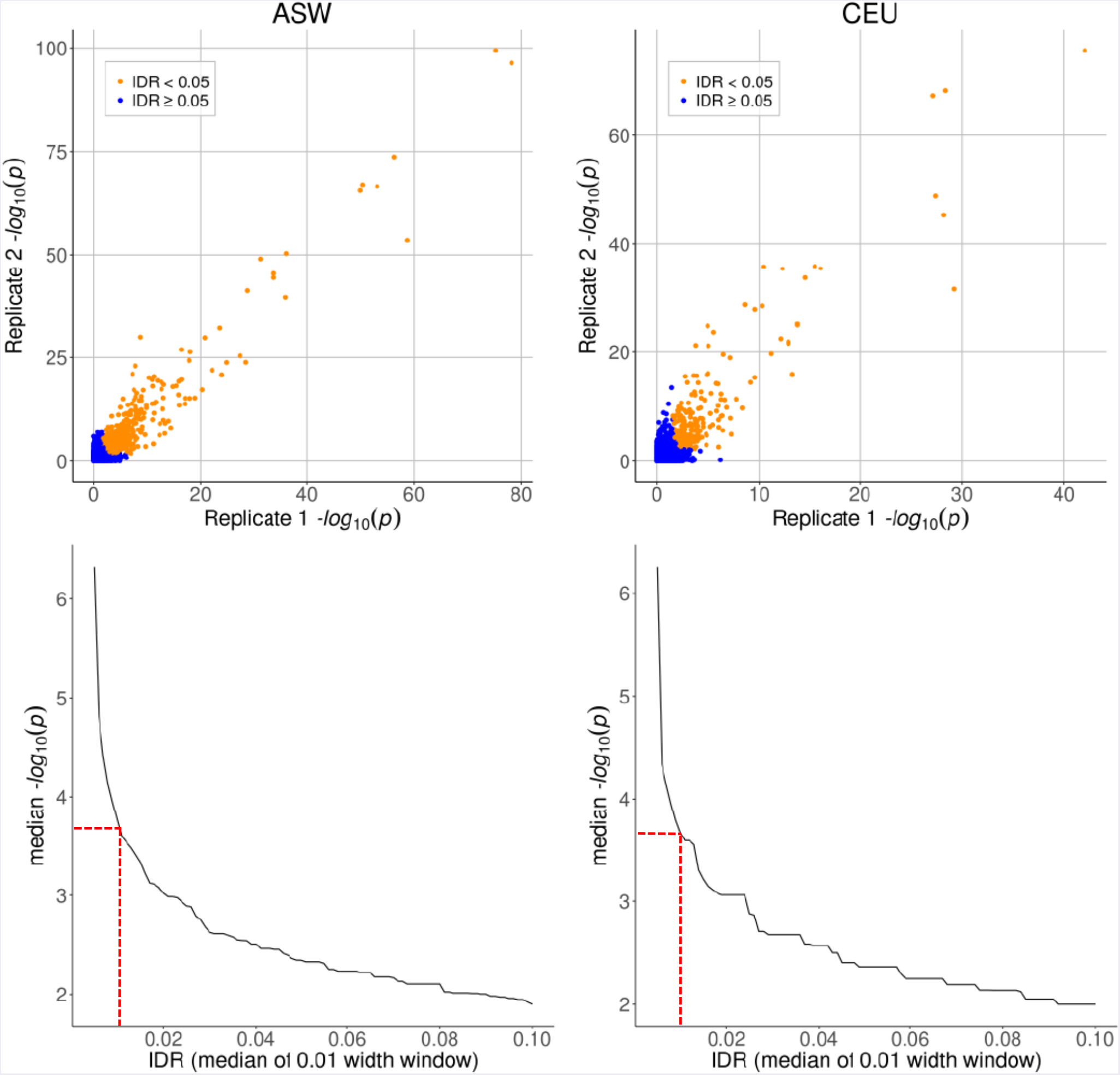
Top row: caQTL p-values for SNPs on chr 1 in ASW and CEU, shown separately for each biological replicate. Bottom row: median -logio(p-value) as a function of IDR, plotted using a moving window of IDR values (window width = 0.01). Dashed red lines indicate the p-value cutoff of 5xl0’^4^, corresponding to IDR ≈ 0.01.

**Figure 1—figure supplement 4.**
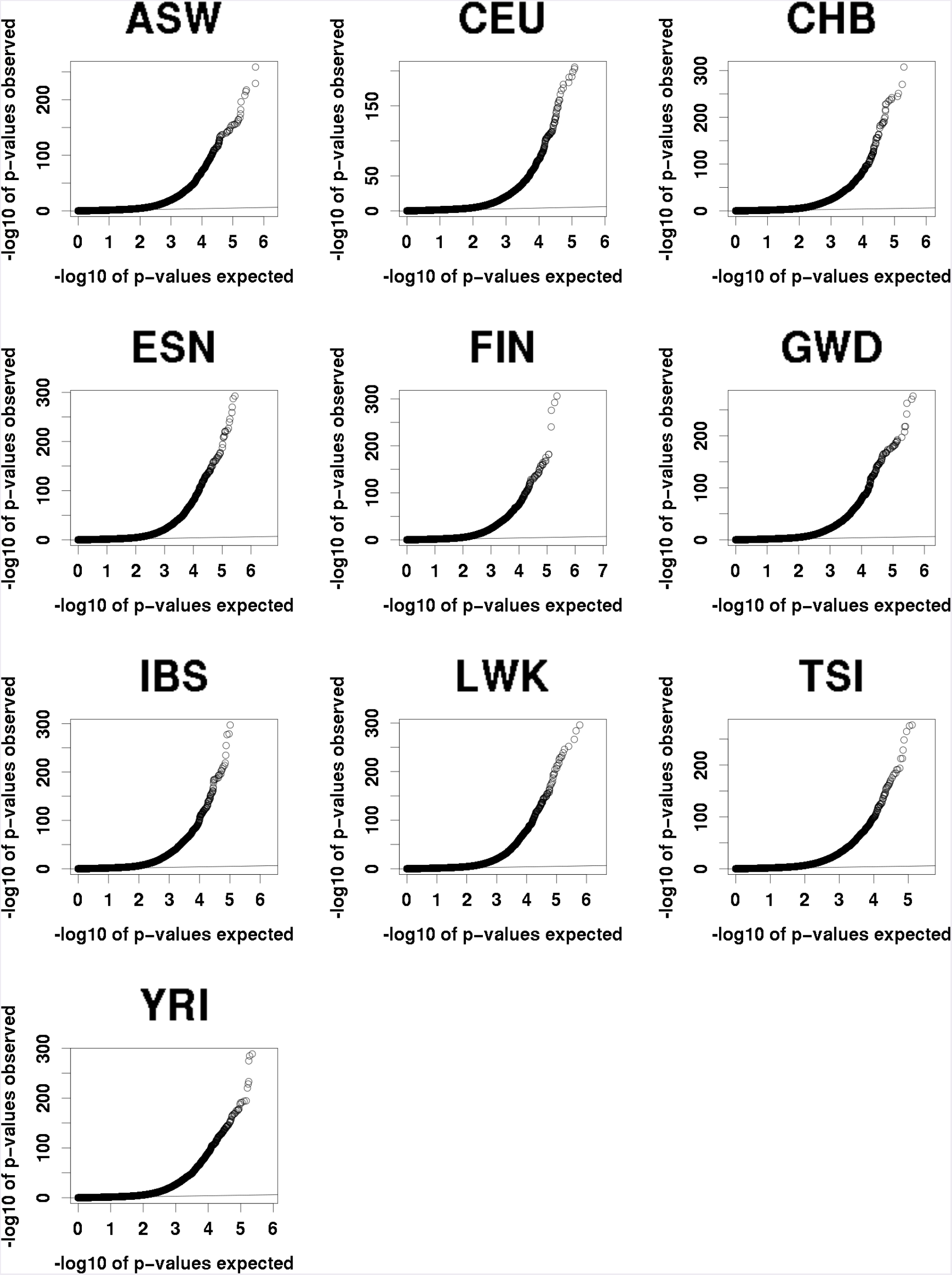
QQ plots of expected (under the null) vs observed caQTLs p-values. All populations show a similar excess of significant p-values.

**Figure 1—figure supplement 5.**
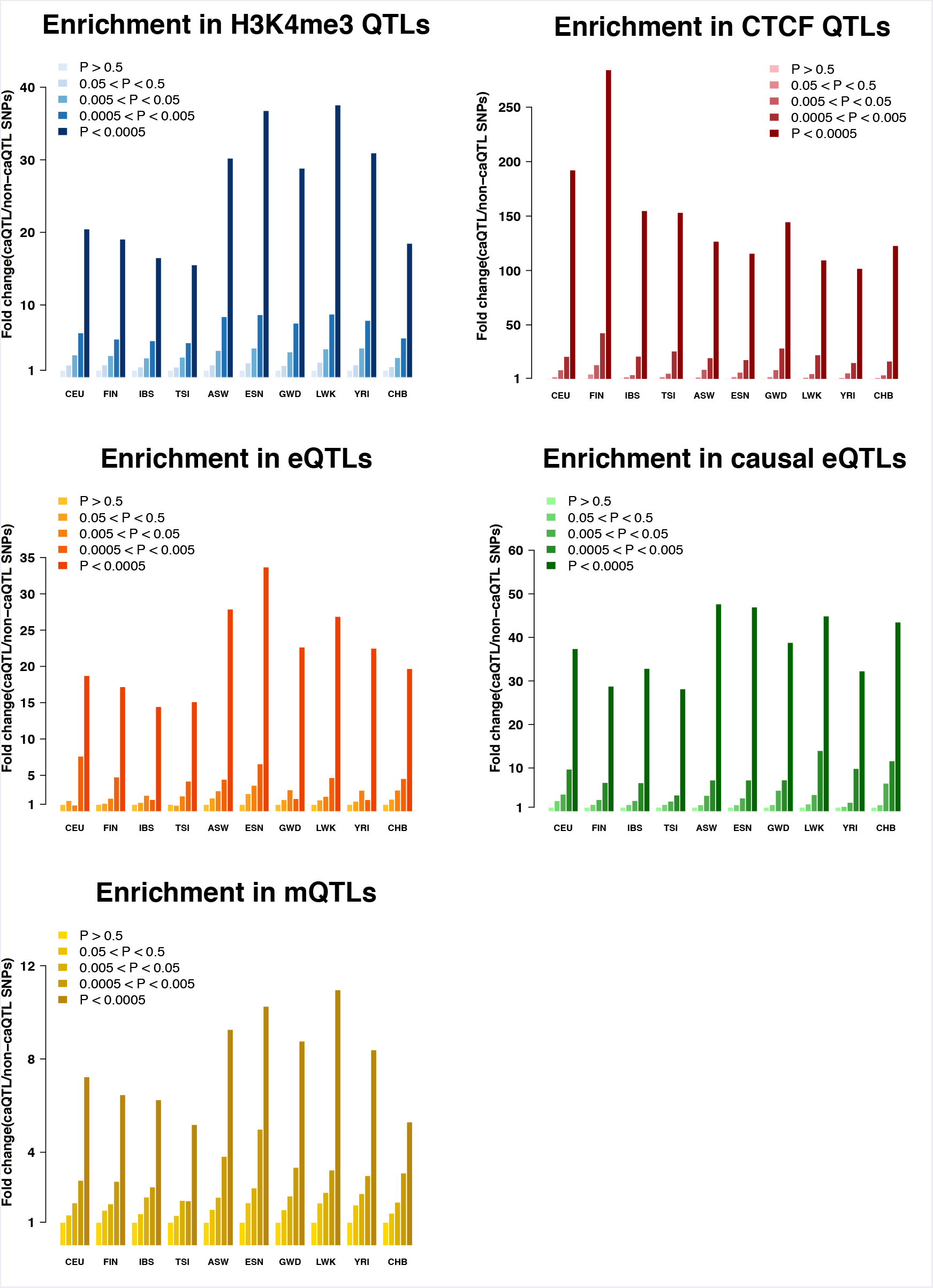
caQTLs enrichments among other molecular QTLs (Ding et al. 2014, Lappalainen et al. 2013, Waszak et al. 2015, Tewhey et al. 2016, Banovich et al. 2014).

**Figure 2—figure supplement 1.**
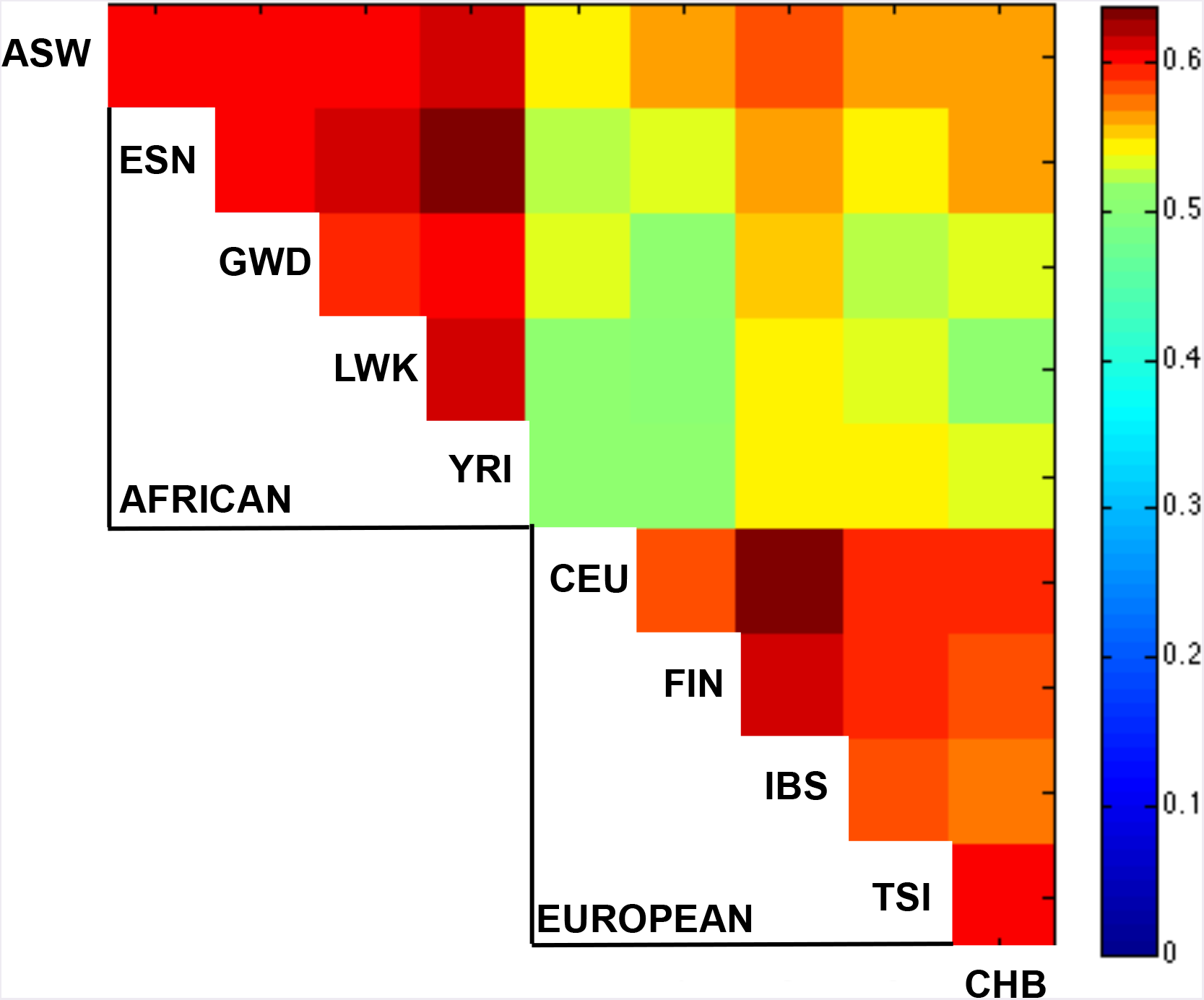
Sharing of caQTLs across populations, as in Fig. 2A, but excluding comparisons with divergent allele frequencies. One possible explanation for the increase sharing of caQTLs between closely related population (Fig. 2A) is that since the allele frequency can affect power to detect QTLs, more similar allele frequencies could lead to greater levels of sharing. To test this possibility, for each SNP, we calculated the sharing as in Fig. 2A after excluding any population that had a pre-ATAC allele frequency >5% away from the mean frequency across all 10 populations. Although this excluded 75% of pairwise comparisons, we still observed a similar pattern of sharing, suggesting that patterns of sharing are unlikely to be driven solely by allele frequency differences.

**Figure 2—figure supplement 2.**
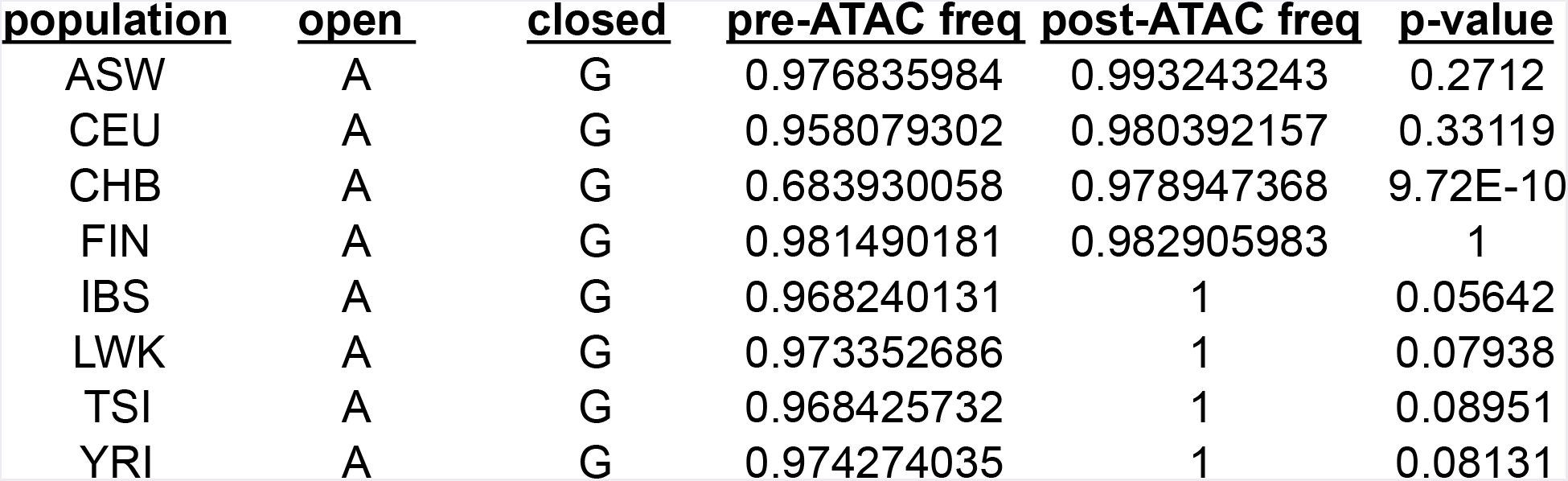
Example of a shared caQTL (rs79979970) that is individually significant in only one population (CHB) out of eight tested, but reaches a shared caQTL p = 5.6x10^-7^ because it has p < 0.1 in an additional four populations. In this case, CHB had the greatest power to detect an effect since it had a pre-ATAC allele frequency of 0.68 for the open allele, whereas the other seven all had frequencies > 0.95 and thus very little range for the open allele to increase in frequency post-ATAC.

**Figure 4—figure supplement 1.**
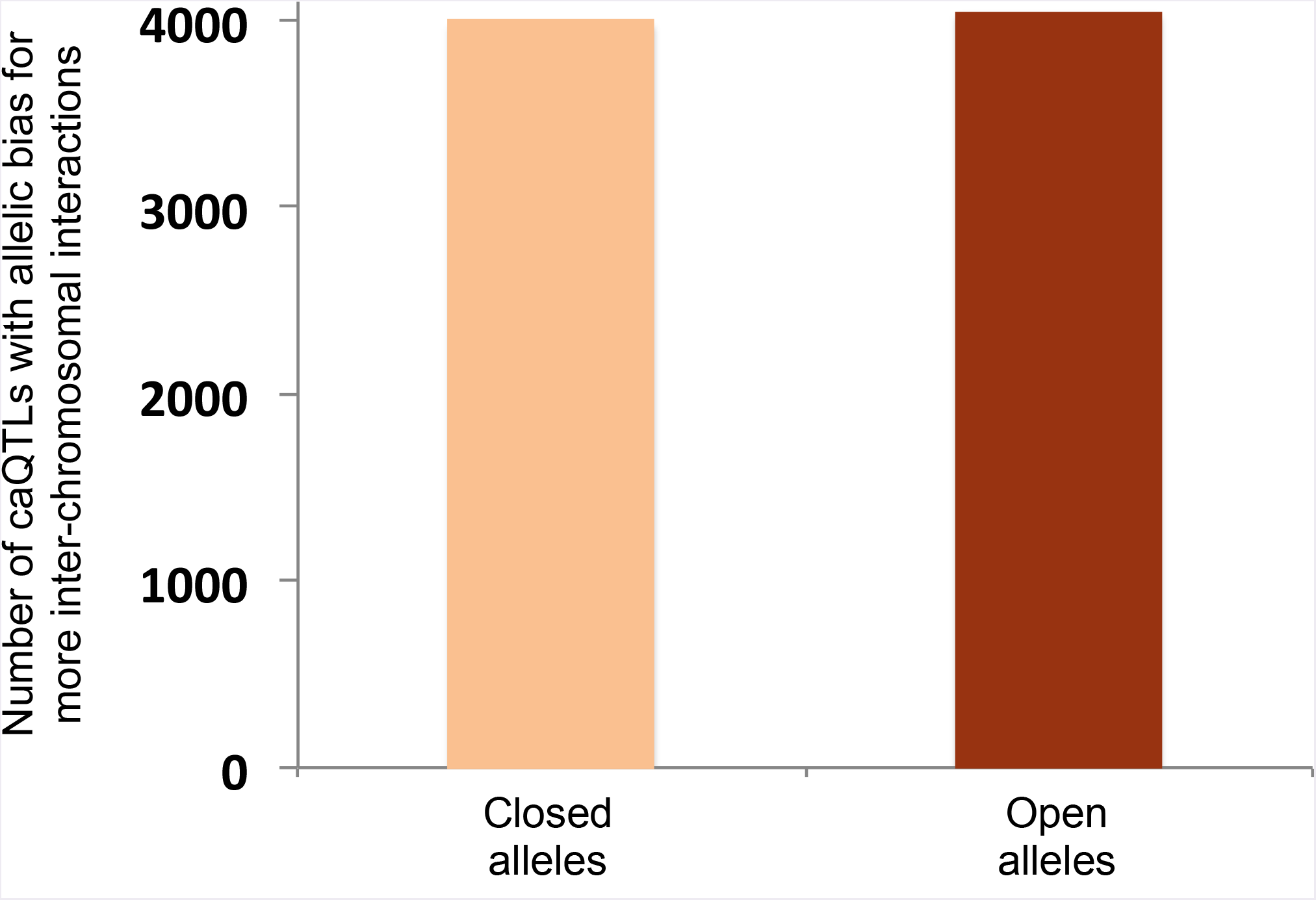
Allelic bias of shared caQTLs for inter-chromosomal interactions. To test the possibility that our result in Fig. 4A could be due to a nonspecific bias in the Hi-C method—such as open chromatin alleles having higher efficiency of shearing, ligation, or some other step—we reasoned that any such bias should also be reflected in the pattern of allele-specific inter-chromosomal interactions (such inter-chromosomal interactions are typically considered to be “noise”, but should still be affected by any nonspecific biases in the method, making them an ideal control). Using the same Hi-C data (Rao et al. 2014), we found only three caQTLs with significant allelic bias in inter-chromosomal reads (2 favoring open alleles and 1 favoring the closed allele at Bonferroni-corrected p<0.05). Moreover, plotting all shared caQTLs with allele-specific Hi-C data from GM12878 (Rao et al. 2014), shown in this figure, we observed no significant difference (4041 caQTLs favoring open alleles vs. 3990 favoring closed alleles; binomial p=0.58).

**Figure 4—figure supplement 2.**
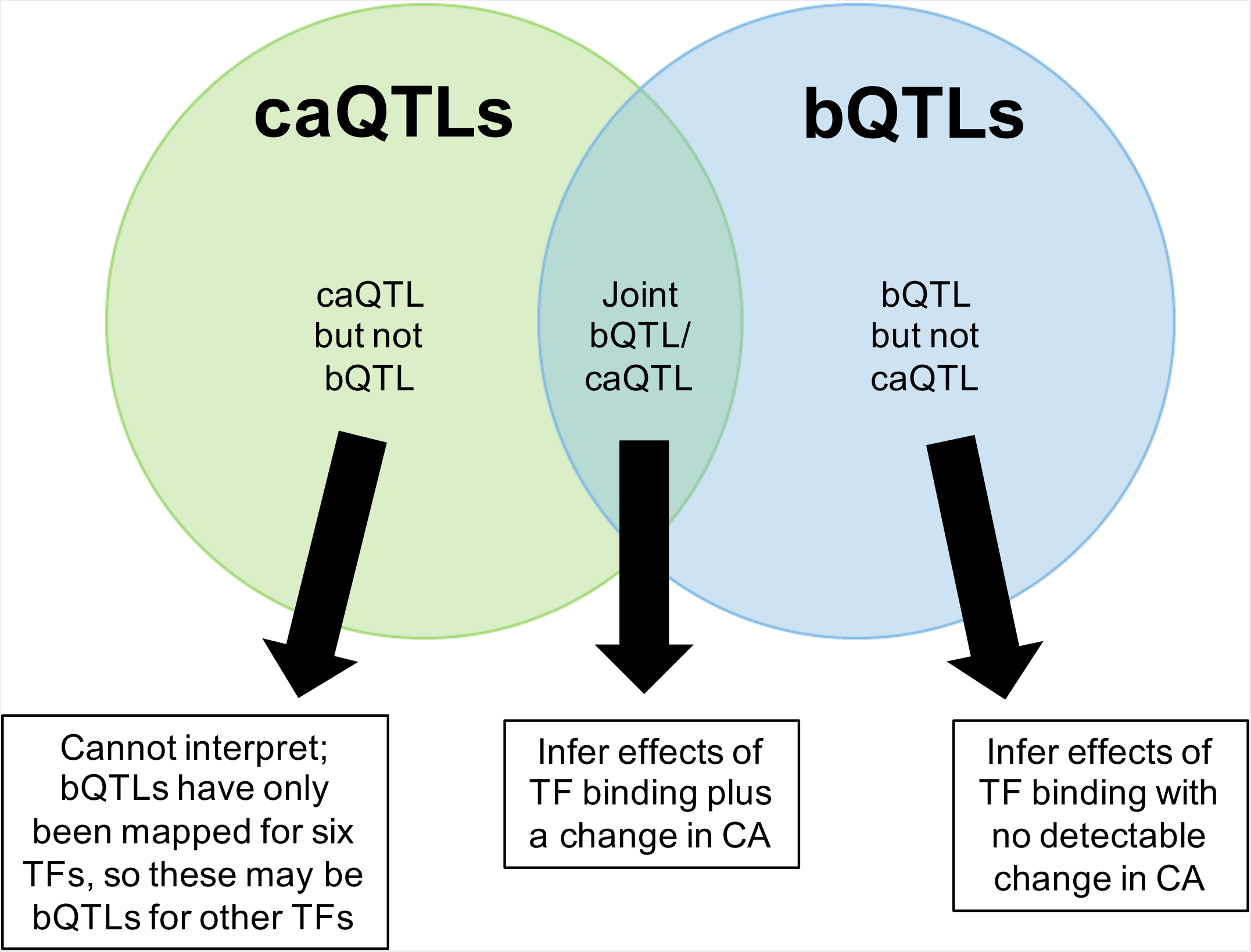
Venn diagram indicating three possible combinations of caQTL/bQTL overlaps, and how we used these to infer their downstream effects in Fig. 4B.

**Figure 4—figure supplement 3.**
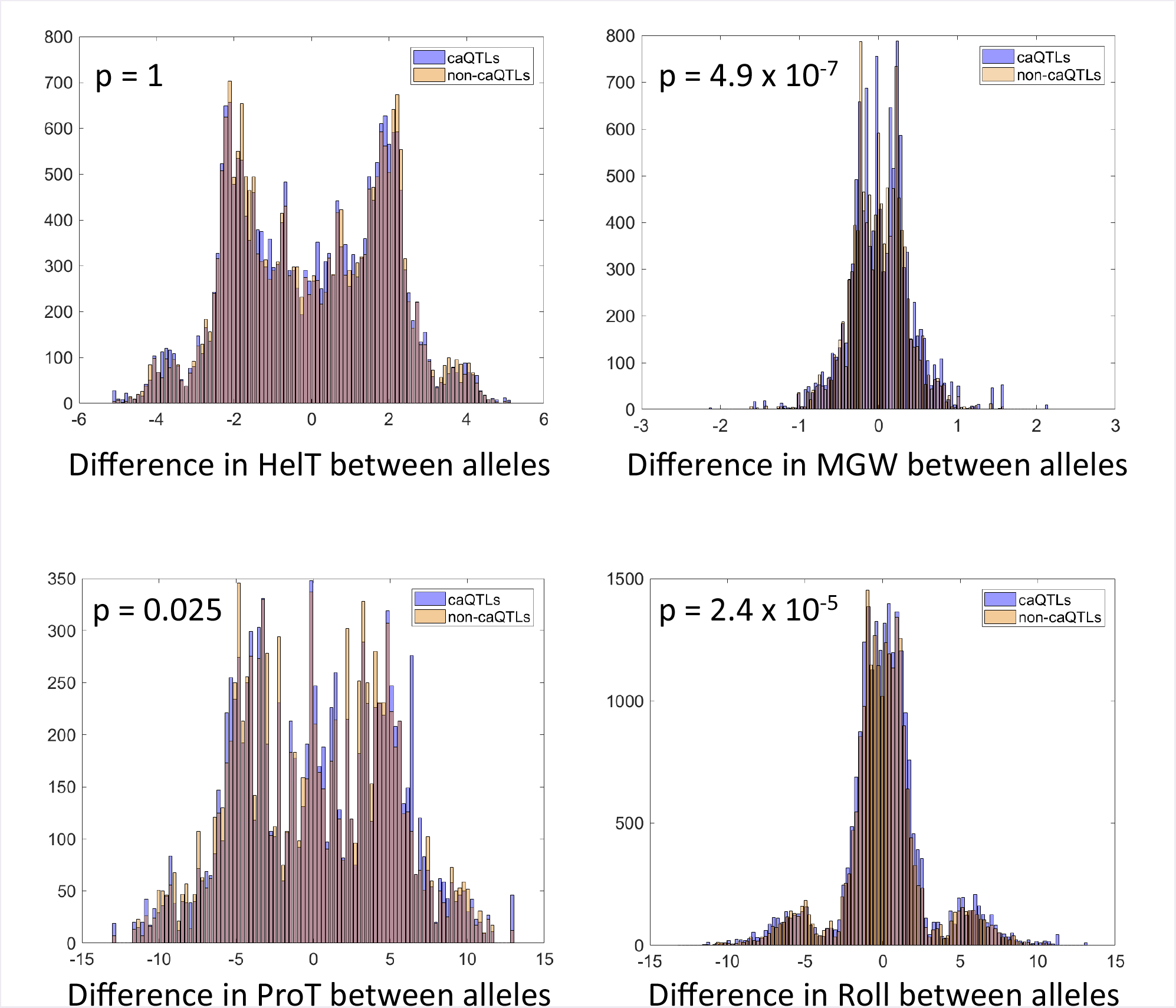
Effect of shared caQTLs on DNA shape. P-values are Bonferroni-corrected for four tests. See Supplemental Note for details.

**Table 1—table supplement 1.**
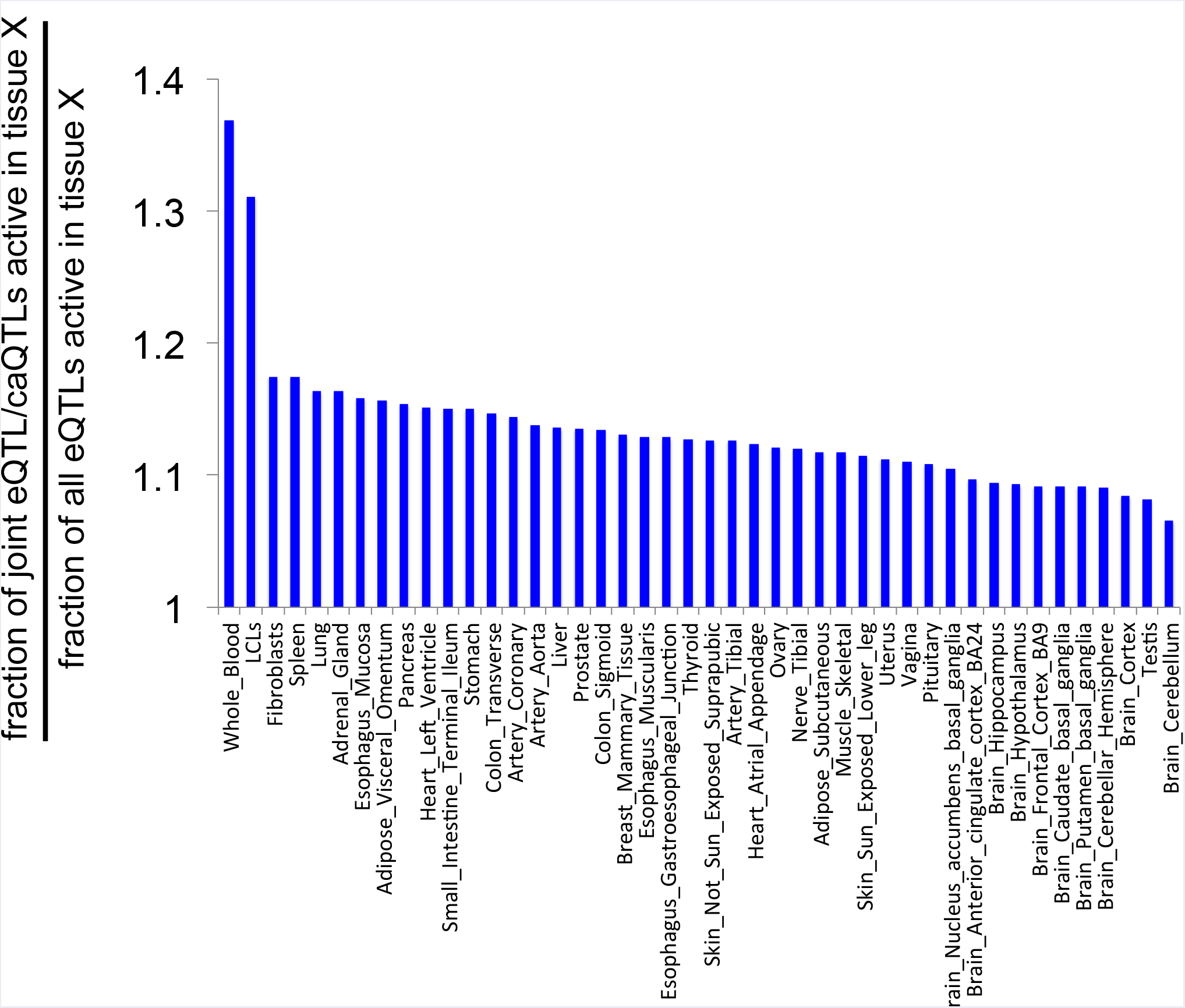
Table 1—table supplement 1. Likelihood of caQTLs from LCLs acting as eQTLs in other tissues. See Supplemental Note for details.

